# Multi-gene biomarkers reveal spatial organization and subpopulation-specific damage response in intrahepatic biliary epithelial cells

**DOI:** 10.64898/2026.02.12.705355

**Authors:** Kendall G Kanakanui, Fransky Hantelys, Hannah R Hrncir, Sergei Bombin, Adam D Gracz

## Abstract

**Background & Aims:** Intrahepatic biliary epithelial cell (BEC) heterogeneity remains challenging to define. Here, we sought to identify BEC subpopulations and biomarkers in mouse liver.

**Methods:** We performed scRNA-seq on Sox9EGFP+ liver epithelium from mice subjected to bile duct ligation (BDL) and sham controls. A machine learning algorithm, NSForest, identified minimal, multi-gene signatures for BEC subpopulations. These “metagenes” were validated using hybridization chain reaction (HCR) FISH in tissue sections from wild-type mice and on primary BECs expanded *in vitro*. Metagenes were used to match BDL subpopulations to their corresponding sham subpopulations for differential gene expression (DGE) analysis.

**Results:** We identified 4 BEC subpopulations in sham controls, each associated with 1-2 gene metagenes. Spatial localization of metagene-defined BEC subpopulations by HCR FISH revealed heterogeneous cellular composition of intrahepatic bile ducts. BECs belonging to a given subpopulation were most likely to have neighbors of the same identity, forming homogenous cellular compartments within ducts. BDL downregulated subpopulation-specific genes and upregulated a damage-associated gene set. BDL samples also included a proliferative subpopulation not found in sham controls, which contained populations enriched for three of the four metagenes. All BEC subpopulations were also found in monolayers *in vitro*, where they clustered spatially with BECs of the same subtype.

**Conclusions:** Novel metagene biomarkers of BEC subpopulations facilitated spatial localization of BECs *in situ*, identified subpopulation specific injury responses, and confirmed that BEC heterogeneity is preserved *in vitro*. The presence of locally homogenous BEC “neighborhoods” *in vitro* suggests some degree of BEC organization may be epithelial-autonomous.

## INTRODUCTION

Biliary epithelial cells (BECs) line the intrahepatic and extrahepatic bile ducts (IHDBs and EHBDs, respectively) forming a continuous network that modifies and transports bile from the liver to the digestive tract. Bile serves critical functions to break down dietary fats for nutrient absorption and excrete waste products like excess cholesterol and xenobiotics. While BECs make up only 3-5% in the healthy liver by cell number, liver injury can induce a proliferative expansion of BECs, termed ductular reaction (DR). DR is thought to have both protective and maladaptive consequences, maintaining bile flow while also resulting in pro-fibrotic and pro-inflammatory signaling ^1, 2^. Overall, the vital role BECs play in regulating bile composition and bile flow is essential for proper liver function and to maintain organismal homeostasis ^3^.

Despite the central role of BECs in liver biology and evidence that subpopulations of BECs exhibit heterogeneous function in homeostasis and response to injury, BEC heterogeneity has remained challenging to define and assay. Classical models have defined BEC subpopulations relative to their localization in small ductules or large ducts ^4-6^. However, many of the biomarkers associated with small and large BECs are either not detectable or broadly expressed in recent transcriptomic data sets ^7-10^. Single cell RNA-seq (scRNA-seq) has facilitated unbiased characterization of BEC heterogeneity but has yet to result in a definitive “ground truth” understanding of BEC subpopulations consisting of broadly accepted cell types and their associated biomarkers. BECs are a relatively rare cell type in the liver and exhibit high cell-cell transcriptomic similarity to one another, resulting in underrepresentation in liver atlases and a lack of distinct “clusters” in unbiased scRNA-seq analyses ^9, 11-13^ . BEC subpopulations also lack well-defined, cell-type specific biomarkers, limiting *a priori* approaches to scRNA-seq analysis and high confidence identification of cell-cell heterogeneity. Prior studies have addressed this challenge by defining BEC subpopulations relative to proliferative status or in terms of gradients of gene expression ^10-13^. For example, Yap signaling was observed to be active in all BECs but expressed at high and low levels in distinct BEC subpopulations ^11^. However, a reliance on transcriptome-level data limits consideration of BEC heterogeneity and subpopulation-specific responses to scRNA-seq assays.

Our lab previously characterized BEC heterogeneity relative to expression of the transcription factor *Sox9*, using a Sox9^EGFP^ BAC transgenic reporter mouse ^8^. Sox9^EGFP^ is expressed in all BECs, but at distinct levels in BECs (termed GFP^low^ and GFP^high^), as well as a rare population of peribiliary hybrid hepatocytes (HybHeps, GFP^sublow^) ^8, 14^. While the Sox9^EGFP^ allele is advantageous as a single biomarker approach for identifying transcriptomically distinct BEC subpopulations, whether it captures the full extent of BEC heterogeneity is unclear. In the present study, we sought to profile BEC heterogeneity and identify experimentally tractable biomarkers of BEC subpopulations by scRNA-seq.

## RESULTS

### Single cell transcriptomic profiling of BECs and hybrid hepatocytes

To produce a single cell atlas focused on BECs, we performed scRNA-seq on EGFP+ cells from both healthy and cholestatic Sox9^EGFP^ livers. Mice were either subjected to bile duct ligation (BDL) or sham surgery and sacrificed after 7 days. We confirmed that EGFP marks BECs (EGFP+/CD326+) and HybHeps (EGFP+/CD326-) in both sham and BDL livers by immunofluorescence, as previously reported (Figure 1A) ^8^. EGFP+ cells were isolated by fluorescence-activated cell sorting (FACS) and subjected to scRNA-seq using split-pool chemistry (Supplemental Figure 1A). Cells with fewer than 100 genes detected or with mitochondrial RNA content higher than 50% were excluded (Supplemental Figure 1B). Following QC filtering, our final dataset contained 6,926 Sox9^EGFP^+ cells from two sham mice and 5,282 Sox9^EGFP^+ cells from two BDL mice. Because no established ground truth exists regarding the number of mouse BEC subpopulations, data were analyzed for optimal cluster number using clustree (Supplemental Figure 2C-D) ^15^. We excluded a cluster of contaminating endothelial cells, marked by high *Pecam1* expression, as well as clusters containing fewer than 100 cells (Supplemental Figure 2E). We identified 13 high confidence clusters across sham and BDL samples (Figure 1B). Interestingly, sham and BDL samples clustered separately from one another, implying that cholestasis has a greater effect than subpopulation identity on BEC transcriptomic heterogeneity. Individual samples were uniformly distributed throughout sham and BDL clusters, suggesting that clustering was not driven by variation between biological replicates (Supplemental Figure 2F).

**Figure 1.**
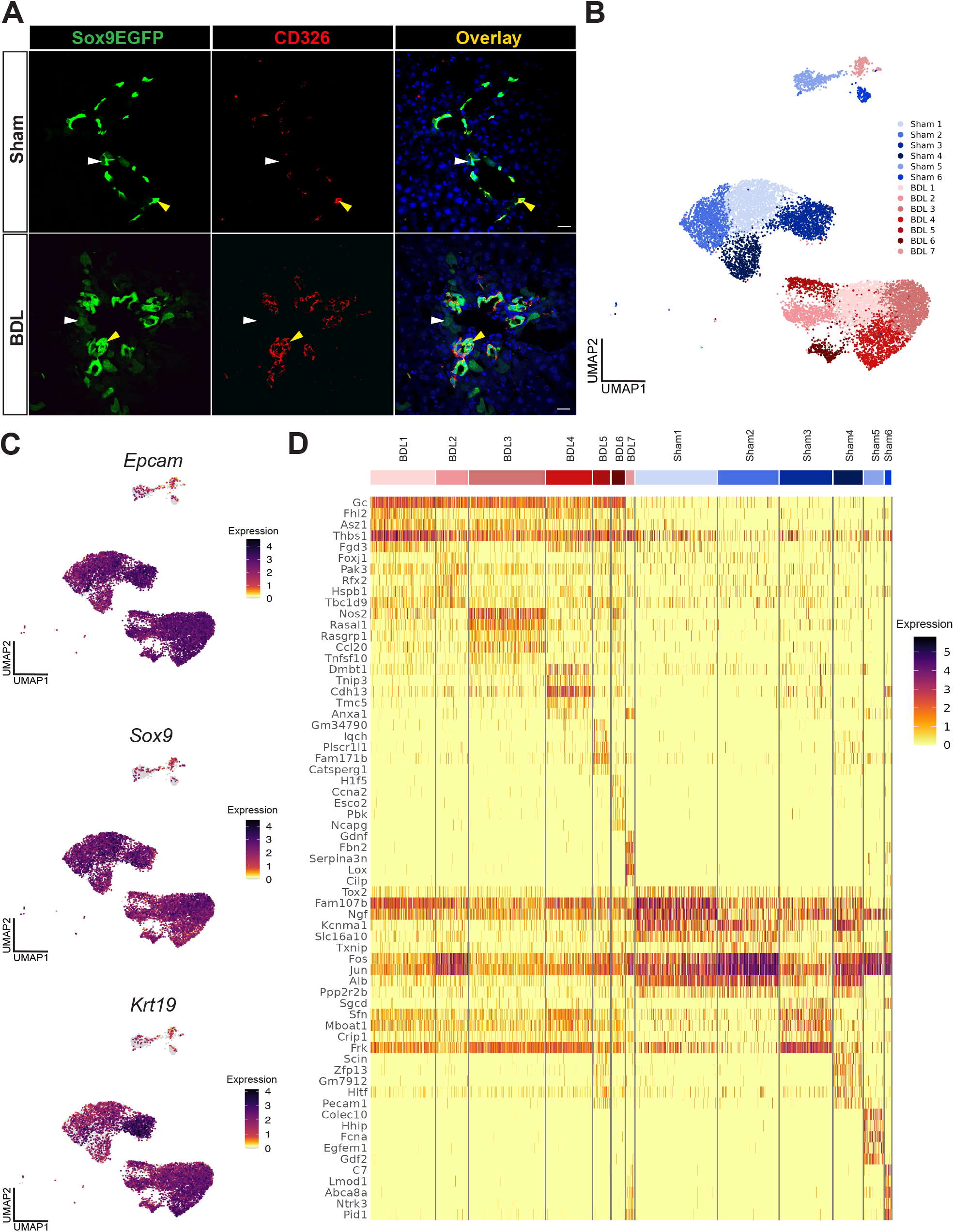
scRNA-seq of Sox9^EGFP^+ cells captures biliary epithelial cells and hybrid hepatocytes in healthy and cholestatic livers. (A) Immunofluorescence of sham operated and 7d post-BDL Sox9^EGFP^ liver confirms that EGFP localizes to CD326+ BECs (marked by white arrowheads) and CD326-periportal HybHeps (marked by yellow arrowheads). (B) EGFP+ cells can be classified into 13 transcriptomically distinct clusters, with differential clustering associated with sham vs. BDL experimental groups. (C) Expression of canonical BEC markers *Epcam, Sox9*, and *Krt19* confirms identity of BEC clusters (Sham 1-4, BDL 1-6). (D) The top 5 differentially expressed genes in BEC clusters (Sham 1-4, BDL 1-6) and HybHep clusters (Sham 5-6, BDL 7) exhibit varying levels of transcriptional heterogeneity across cell type and experimental group. Top genes in sham BEC clusters are broadly expressed across clusters, suggesting substantial transcriptomic similarity between individual BECs.

Since Sox9^EGFP^ is expressed in both BECs and HybHeps, we next sought to confirm epithelial lineage identity of scRNA-seq clusters. First, we examined *Egfp*, which is known to be expressed highly in BECs and at low levels in HybHeps ^8^. *Egfp* was enriched in clusters Sham 1-4 and BDL 1-6, suggesting that these clusters represent BECs while Sham 5, Sham 6, and BDL 7 represent HybHeps (Supplemental Figure 2A). To further confirm BEC and HybHep identity of clusters, we scored transcriptomic signatures of the three Sox9^EGFP^ populations (GFP^high^, GFP^low^, and GFP^sublow^) derived from published bulk RNA-seq ^8^. GFP^high^ and GFP^low^ signatures, which are associated with BEC populations, were enriched in Sham 1-4 and BDL 1-6, consistent with *Egfp* expression in these clusters (Supplemental Figure 2C & D). The HybHep GFP^sub^ gene signature was enriched in Sham 5, Sham 6, and BDL 7 (Supplemental Figure 2B). Together, these data identify BEC (Sham 1-4 and BDL 1-6) and HybHep (Sham 5 & 6, BDL 7) clusters in our larger dataset.

Previous studies applying scRNA-seq to BECs note: (1) broad expression of universal BEC biomarkers across computationally identified clusters and (2) subpopulation-specific expression of genes associated with hepatocytes ^7, 10, 12^. To examine this in our own dataset, we analyzed expression patterns of canonical BEC genes (*Cd326, Krt19*, and *Sox9*) and canonical hepatocyte genes (*Hnf4a* and *Alb*). As expected, BEC genes were broadly expressed across BEC clusters and showed lower expression in HybHep clusters (Figure 1C). *Krt19* was further enriched in Sham 3, consistent with prior observations that the Sox9^EGFP-low^ subpopulation expresses elevated *Krt19* relative to other BECs (Figure 1C & Supplemental Figure 2B) ^8^. Unexpectedly, *Hnf4a* and *Alb* exhibited higher expression in BEC clusters than in HybHep clusters (Supplemental Figure 3A). Hepatocyte biomarker expression was also somewhat variable between BEC clusters, with decreased *Hnf4a* expression in Sham 2 and 4 and decreased *Alb* expression in BDL 3 relative to other BEC clusters (Supplemental Figure 3A). We also examined expression of reported small ductule and large duct markers, which represent classical markers of BEC heterogeneity^4, 6, 16-18^. While the expression of small BEC marker *Hrh1* was enriched in Sham 3 and BDL 3-4, other markers were either broadly expressed in BEC clusters or exhibited rare, scattered detection across clusters (Supplemental Figure 3B & C). Therefore, expression of canonical BEC and hepatocyte genes reinforces the overall epithelial identity of clusters in our datasets and confirms previously noted transcriptomic similarities between BECs.

Prior studies have relied on enrichment of gene sets associated with proliferation, hepatocyte identity, or Yap signaling to define differences between otherwise transcriptomically similar BEC subpopulations ^11, 12^. To test if a similar approach could discriminate clusters in our dataset, we curated eight gene sets related to BEC specification and function from published data, including gene sets discriminating large ducts from small ductules ^7, 19-21^. Gene sets for Yap, Notch, and primary cilia were enriched in BEC clusters, whereas Hedgehog and ECM gene sets were enriched in HybHep clusters (Supplemental Figure 2E-I). Yap, Notch, and TGF-β signaling were generally upregulated in BDL clusters relative to Sham clusters, supporting injury-induced activation of these pathways (Supplemental Figure 2E-F, 2J) ^12, 22, 23^. TGF-β was most enriched in BDL 1 and BDL 3, indicating possible subpopulation-specific activation (Supplemental Figure 2J). Additionally, genes upregulated in large ducts vs small ductules were enriched in Sham 3 and BDL 3, suggesting that BEC identity associated with anatomical location in the IHBDs is maintained in both homeostasis and injury (Supplemental Figure 2K). While gene set enrichment analyses (GSEA) reveal subpopulation-specific differences that reinforce differences between BEC subpopulations, they mainly exhibit subtle relative enrichment or de-enrichment across BEC clusters. Further, because the gene sets examined here consist of 23-740 genes, they remain impractical as biomarkers of BEC subpopulations in assays other than scRNA-seq.

To pursue unbiased identification of individual genes that could serve as BEC subpopulation biomarkers, we performed differential gene expression between all clusters (Supplemental Table 1). The top 5 differentially expressed genes (DEGs) from each cluster exhibited distinct expression between: (1) BEC and HybHep populations and (2) sham and BDL populations (Figure 1D). However, like previously examined gene sets associated with BECs, top DEGs for each BEC cluster exhibited either: (1) low coverage of individual cells within its cluster or (2) broad expression across other BEC clusters (Figure 1D). Together, these findings demonstrate substantial transcriptomic differences between BECs and HybHeps and BECs from healthy livers vs. BECs from livers subjected to BDL. As with other published scRNA-seq data sets, our analyses also suggest that BEC subpopulations are highly similar at the transcriptomic level, complicating identification of subpopulation-specific biomarkers.

### Candidate metagenes define BEC subpopulations in homeostasis

To identify “ground truth” biomarkers for BEC subpopulations, we next focused only on cells from sham controls and performed differential gene expression analysis to identify candidate biomarkers during BEC homeostasis (Supplemental Table 2). We identified 1,853 DEGs, with Sham 1- and Sham 2-associated DEGs exhibiting broad expression across all four clusters while Sham 3- and Sham 4-associated DEGs were more restricted to their respective clusters (Figure 2A). GSEA revealed subpopulation-specific enrichment in gene sets consistent with BEC and/or epithelial function (Figure 2B). Importantly, GSEA results for each cluster exhibited non-overlapping significant hits, supporting the unique identity of the four putative BEC subpopulations in our data set.

**Figure 2.**
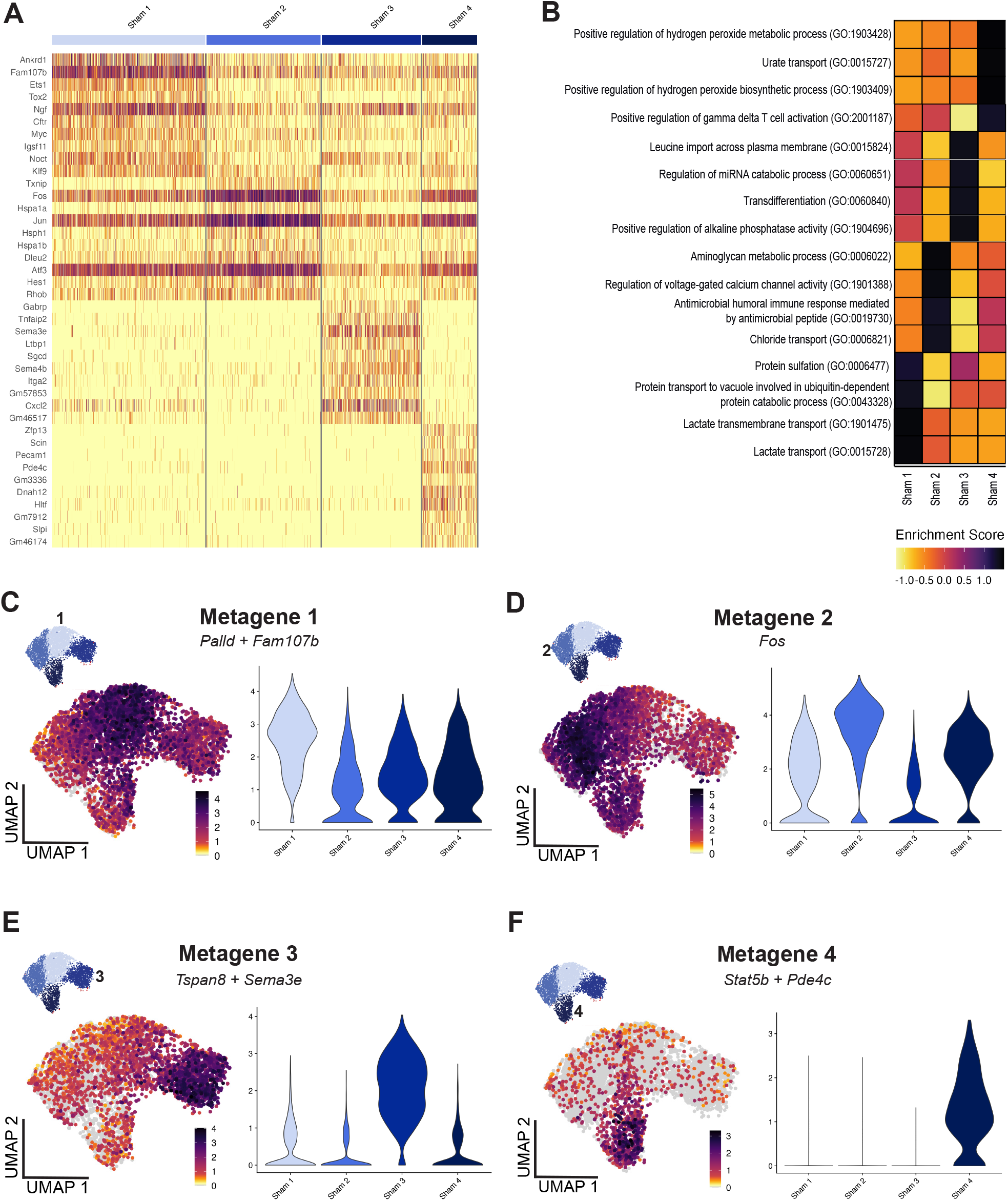
BEC subpopulations in sham operated controls can be defined by the expression of “metagene” biomarkers. (A) The top 10 differentially expressed genes in Sham 1-4 support high transcriptomic similarities between BECs, based on broadly expressed genes. (B) GSEA suggests that sham BEC subpopulations exhibit different functional properties, supporting the unique identity of each cluster. (C-F) Differential expression of candidate metagenes across the four sham BEC subpopulations supports their potential as biomarkers of BEC heterogeneity. Each metagene demonstrates enrichment in its “on target” cluster is de-enrichment in “off target” clusters.

DEGs identified for Sham 3 and Sham 4 exhibited “binary” expression, an important characteristic of potential biomarkers defined by enrichment in the “target” cluster and simultaneous de-enrichment in all other clusters ^24^ . However, DEGs identified for Sham 1 and Sham 2 did not exhibit binary expression, as many were also expressed at variable levels in Sham 3 and Sham 4 (Figure 2A). To identify genes with binary expression that might not be represented in the top DEGs for each cluster, we applied NSForest, a previously published machine learning algorithm ^24, 25^. NSForest identifies potential biomarkers in scRNA-seq data by scoring genes for both binary expression as well as expression in all cells within a defined cluster ^24^. We identified four minimal gene biomarkers composed of 1-2 genes associated with each subpopulation, which we termed “metagenes” (Supplemental Figure 4). *Palld* and *Fam107b* (Metagene 1) identify Sham 1, *Fos* (Metagene 2) defines Sham 2, *Tspan8* and *Sema3e* (Metagene 3) define Sham 3, and *Stat5b* and *Pde4* (Metagene 4) define Sham 4 (Figure 2C-F). *Palld* encodes a cytoskeletal protein involved in actin organization and *Fam107b* encodes a nuclear protein likely involved in stress response ^26, 27^. Metagene 2 consists only of *Fos*, which encodes a subunit of the transcription factor AP-1 and is associated with hepatocellular carcinoma (HCC) in humans ^28^. In Metagene 3, *Tspan8* encodes a transmembrane protein important in lipid metabolism and *Sema3e* encodes a secreted class 3 semaphorin involved in several context-dependent processes, such as its regulation of sinusoidal regeneration in liver fibrosis during liver regeneration ^29-32^. Both genes have been identified as markers for several cancers, and specifically *Tspan8* has shown to regulate the stemness of breast and pancreatic cancer cells ^12, 33-35^. In Metagene 4, *Stat5b* encodes a transcription factor with potential fibrotic and metabolic regulatory roles and *Pde4c* encodes an enzyme that functions in cAMP signaling specifically associated with inflammation ^36, 37^. *Tspan8* has been proposed to be a novel BEC marker, but to our knowledge, the genes comprising the metagenes have not been previously proposed as subpopulation-specific biomarkers for BECs ^12^. We next compared the expression of all four metagenes across Sham BEC subpopulations.

Only Metagene 4 exhibited striking binary expression (Figure 2F). The other three metagenes had varying degrees of binary expression, with Metagene 3 demonstrating the next most restrictive expression and Metagene 1 demonstrating the broadest expression across BEC subpopulations (Figure 2C-E). Interestingly, metagenes consisting of two genes (Metagene 1, 3, and 4) have a similar pattern of co-expression. One gene was highly expressed in its specific cluster (*Palld, Tspan8*, and *Stat5b*), but also in other clusters (Supplemental Figure 4A, D, & F). The other gene was more restricted to its specific cluster (*Fam107b, Sema3e*, and *Pde4c*), but exhibited lower coverage of individual cells within its cluster (Supplemental Figure 4B, E, & G). As a panel, these metagenes represent potential biomarkers for identifying BEC subpopulations, which can be identified by: (1) enrichment of the cluster specific metagene and (2) the de-enrichment of metagenes associated with other BEC subpopulations.

### Spatial validation of BEC metagenes identifies homogenous cellular neighborhoods within heterogeneous ducts

To test whether candidate metagenes could function as BEC subpopulation biomarkers *in situ*, we designed a 7 gene panel for hybridization chain reaction (HCR) FISH and applied this to liver tissue sections from wild-type C57Bl/6 mice (Figure 3A) ^38^. Because HCR FISH allows for relative quantification of gene expression, BECs were segmented based on EPCAM signal and scored based on relative metagene expression to derive a subpopulation assignment (see Methods; Figure 3B and Supplemental Table 3). Each BEC demonstrated clear enrichment of a single metagene and de-enrichment of the other three metagenes, as observed in scRNA-seq analysis (Supplemental Table 3). We quantified 243 BECs across 23 fields from three biological replicates, and identified 24.69% of BECs enriched for Metagene 1, 23.87% of BECs enriched for Metagene 2, 28.81% of BECs enriched for Metagene 3, and 22.63% of BECs enriched for Metagene 4. We compared the proportion of each BEC subpopulation identified in HCR FISH data to proportions from scRNA-seq and observed high concordance between the two detection methods (Figure 3C). Notably, differences in proportions of each BEC subpopulation in HCR FISH and scRNA-seq amounted to less than 12% (Figure 3C). These results suggest that *in situ* quantification of candidate metagenes is sufficient to discern BEC subpopulations identified by scRNA-seq. To simplify discussion of BEC subpopulations throughout the rest of this manuscript, each scRNA-seq identified cluster will be referred to by its corresponding Metagene number (for example, BECs enriched for Metagene 1 are referred to as “BEC-I”).

**Figure 3.**
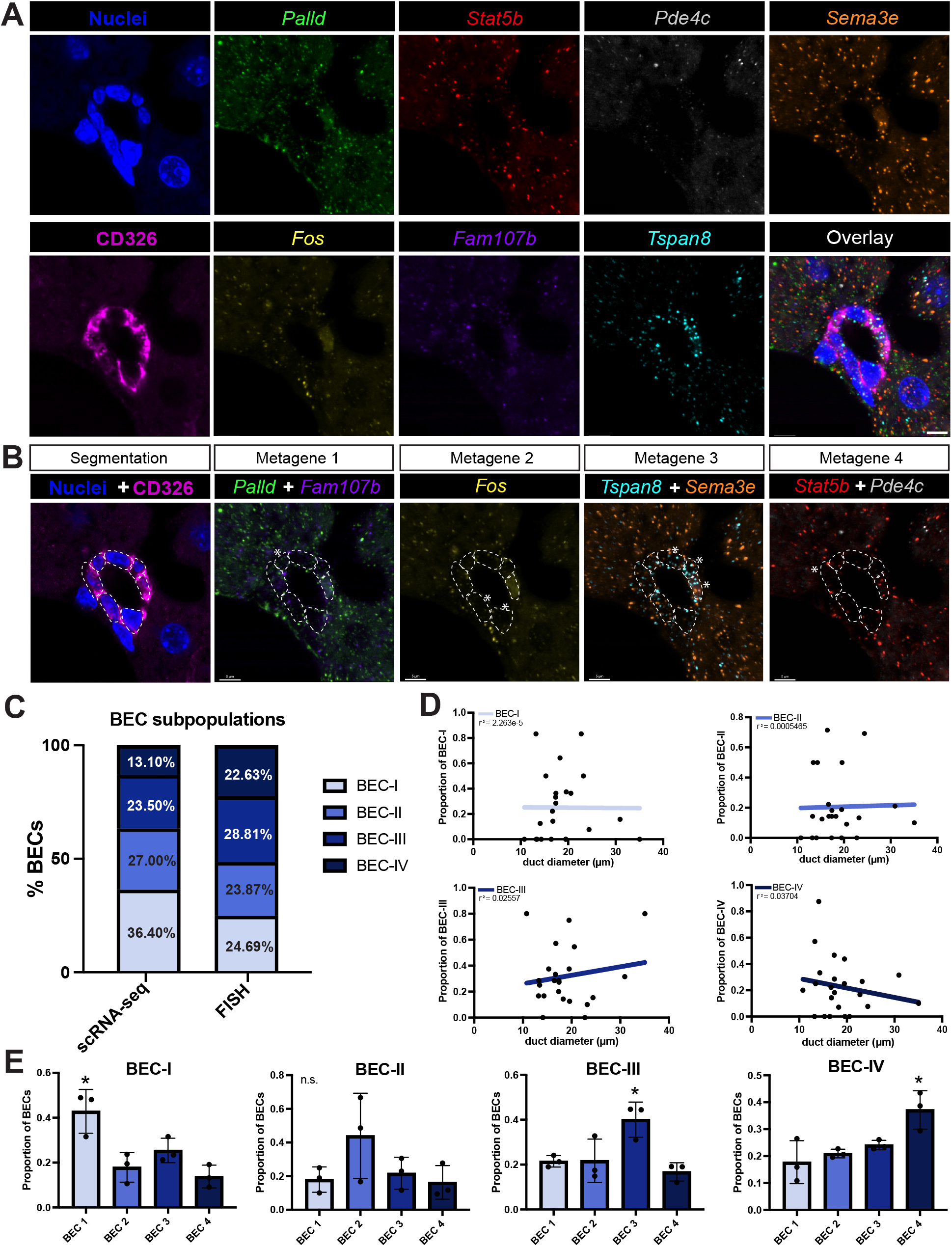
HCR FISH for candidate metagenes validates BEC subpopulations in vivo and provides spatial context for BEC heterogeneity. (A) Validation of a 7-plex HCR FISH panel (*Palld, Stat5b, Pde4c, Sema3e, Fos, Fam107b, Tspan8*) for metagene detection in liver tissue from wild-type, C57Bl/6 mice. (scale bar represents 10µm). (B) BEC identification and segmentation by CD326 facilitates metagene quantification and classification of individual BECs by subpopulation identity (asterisk marks the BEC identity within each metagene-specific image field.) (scale bar represents 10µm). (C) Proportions of each BEC subpopulation identified by HCR FISH are similar to proportions in scRNA-seq data. (D) Quantification of BEC identity relative to IHBD diameter reveals a trend for BEC-III localization to larger ducts and BEC-IV localization to smaller ducts. (E) Quantification of neighboring cell identities reveals BECs are likely to have neighboring BECs of the same subtype (asterisk indicates significance, p<0.05. Data in panels C-H were generated from 3 independent liver samples).

Next, we sought to leverage *in situ* application of the metagene panel to gain insight on the spatial organization of BEC subpopulations within IHBDs. All IHBDs imaged contained at least two subpopulations (Supplemental Figure 5C). Previous studies have suggested that differences in IHBD size based on duct diameter are linked BEC identity ^6^. While correlation between BEC subpopulation and IHBD size did not reach significance, BEC-III exhibited a weak correlation with larger ducts and BEC-IV with smaller ductules (Figure 3D). This result is consistent with our prior studies demonstrating an increased proportion of Sox9^EGFP-low^ BECs in large IHBDs, as both GFP^low^ and BEC-III are enriched for *Krt19* (Figure 1C and Supplemental Figure 2C) ^8^. The enrichment of GFP^low^ and large duct gene signatures in BEC-III vs. BEC-I, -II, and -IV are also supportive of BEC-III being more represented in larger IHBDs (Supplemental Figure 2C). To further examine the spatial organization of BECs, we quantified the subpopulation identity of neighboring cells for each individual BEC. BECs of each subpopulation were most likely to have a neighboring BEC of the same subpopulation, with the exception of BEC-II, which exhibited a similar trend that did not reach significance (Figure 3E). Spatial data derived from HCR FISH collectively demonstrates that BEC subpopulations are distributed throughout IHBDs, with BEC-III enriched in larger ducts. Additionally, these data reveal that BECs are localized to relatively homogenous epithelial “neighborhoods” within heterogeneous IHBDs.

### Metagenes facilitate identification of subpopulation-specific responses to cholestatic injury

A challenge presented by our initial scRNA-seq analysis was the inability to confidently match Sham clusters with their corresponding BDL cluster due to the substantial transcriptomic differences between each experimental group (Figure 1B & 1F). Additionally, BDL BECs exhibited two additional clusters, further complicating the subpopulation identification across experimental condition. Together, these technical challenges hindered our ability to examine BEC subpopulation specific gene expression changes. We reasoned that we could use metagene enrichment to identify Sham and BDL clusters of the same BEC subpopulation identity. We first examined the enrichment of Metagene 1-4 in BDL samples and found that Metagenes 1, 2, and 4 were enriched in a single BDL cluster (Figure 4A). Given the shared enrichment of Metagene 3 in BDL 3 and BDL 4, we merged the two clusters to maintain a conservative clustering strategy and avoid overrepresenting control-defined BEC subpopulations. (Supplemental Figure 6A). To keep our subpopulation nomenclature consistent, we relabeled the BDL clusters to match their metagene number (Figure 3B). The percentage of cells in each subpopulation following BDL shifted in comparison to sham, with an increase in BEC-III (+28.37%) and concomitant decreases in BEC-I (-9.04%), BEC-II (-13.61%), and BEC-IV (-5.72%) (Figure 4C).

**Figure 4.**
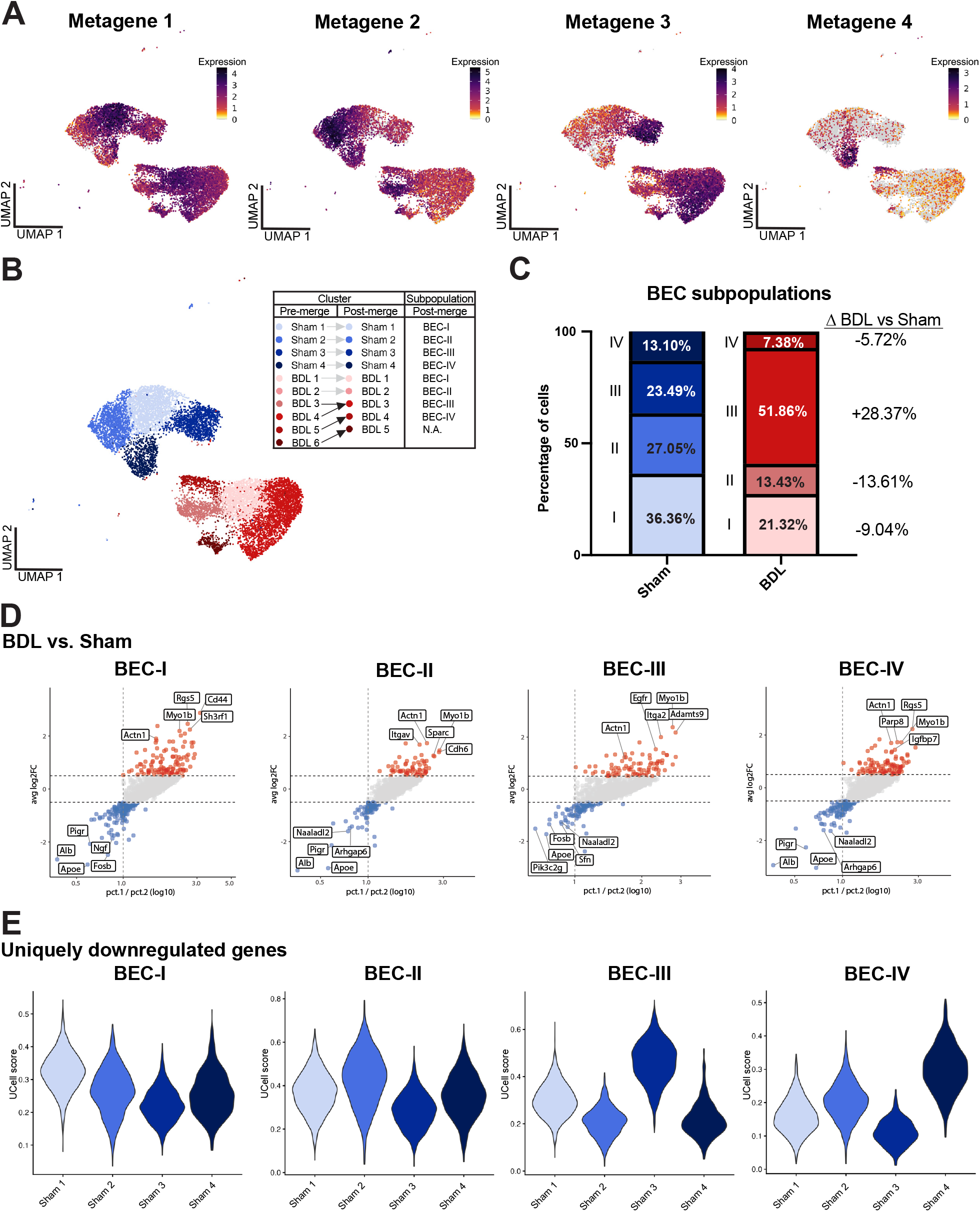
Analysis of metagene-matched sham and BDL clusters reveals shared and subpopulation-specific responses to cholestasis in BECs. (A) Metagene expression facilitates identification of BEC subpopulations in BDL samples that correspond to subpopulations identified in sham controls. Metagene 1 matches BEC-I to BDL 1, metagene 2 matches BEC-II to BDL 2, metagene 3 matches BEC-III to both BDL 3 and BDL 4, and Metagene 4 matches BEC-IV to BDL 5. (B) The BEC clusters were reassigned based on metagene enrichment (Sham 1-4 and BDL 1-5), where BDL 3 and BDL 4 merge into one cluster representing BEC-III, and BDL 5 is relabeled BDL 4 to represent BEC-IV. (C) BDL is associated with a change in BEC proportions by subpopulation, with an increase in BEC-III and corresponding decrease in all other subpopulations. (D) Differential gene expression analysis comparing each BEC subpopulation identifies significant upregulated and downregulated genes in BDL. Dots in red represent significantly upregulated genes (LFC >0.5) and dots in blue represent significantly downregulated genes (LFC <-0.5). The pct1/pct2 ratio is a representation of the effect of BDL on gene expression changes. Notably, genes associated with cell adhesion and cell locomotion are upregulated across all BEC subpopulations (*Actn1* and *Myo1b*), revealing a shared response to cholestasis. (E) Genes downregulated by BDL in a single BEC subpopulation are specifically enriched in that subpopulation in sham controls, suggesting that BDL suppresses subpopulation-specific transcriptomic identity in BECs.

We performed differential gene expression analysis comparing each BDL subpopulation to its respective sham subpopulation, using metagene enrichment as a determinant of BEC identity (Supplemental Tables 4-7). We identified 539 DEGs across all four subpopulation comparisons, of which 293 genes were shared between two or more comparisons (Supplemental Figure 6B). Based on the top 10 upregulated pathways by GSEA, terms related to cell adhesion, locomotion and motility, and cell organization were shared across all pairwise subpopulation comparisons (Supplemental Figure 6C). Examining the top 15 significantly upregulated genes for each subpopulation comparison, two genes were shared across all four comparisons (*Myo1b, Actn1*) (Figure 4D). Both *Myo1b* and *Actn1* have been shown to be major regulators of adhesion and migration ^39, 40^. Together, these data reveal BECs respond globally to cholestasis by upregulating gene expression associated with cellular adhesion and migration. Interestingly, the top 10 downregulated pathways in each of the pairwise comparisons had no terms shared across all four subpopulations (Supplemental Figure 6C). We hypothesized that BDL might be inducing injury-responsive genes broadly across BEC subpopulations while simultaneously repressing subpopulation-specific genes. To test the latter hypothesis, we examined expression of the four sets of uniquely downregulated DEGs in pairwise “Sham vs. BDL” comparisons across all BEC clusters. Because DE analyses of scRNA-seq can be underpowered, we first confirmed that each gene set was uniquely downregulated in its respective BDL cluster, validating differential gene expression analysis (Supplemental Figure 6D). Strikingly, each gene set was also enriched in its respective Sham subpopulation relative to all other Sham subpopulation (Figure 4E). Taken together, these data support a model where BDL induces gene expression programs consistent with cellular adhesion and migration across all BEC subpopulations while simultaneously downregulating subpopulation-specific genes.

In addition to the four metagene-defined BEC subpopulations in BDL, we observed a fifth cluster (BDL 5) marked by robust expression of several cell cycle genes (*Mki67, Cdc20*, and *Cdk1*) (Figure 5A). During homeostasis, approximately 1-5% of BECs are proliferative by KI-67 staining ^41^. Following injury, IHBDs undergo ductular reaction marked by significantly increased BEC proliferation, consistent proportions of *Mki67+* cells in our dataset (Figure 5B). Studies focused on BEC proliferation in response to ductular reaction found that BECs distributed throughout the IHBDs are capable of proliferating, but did not determine whether this response is restricted to a single BEC subpopulation ^42^. Therefore, we sought to test which BEC subpopulations contribute to proliferation by analyzing the expression pattern of the four metagenes in BDL-5. We first subclustered BDL-5 into three clusters (C1-3) based on clustree analysis (Figure 5C & Supplemental Figure 7A). Metagenes 1-3 exhibited enrichment in a single subcluster, while Metagene 4 was reliably expressed in BDL-5 (Figure 5D). These data suggest that BEC I-III are capable of entering the cell cycle following injury and that proliferative responses are not restricted to a single BEC subpopulation.

**Figure 5.**
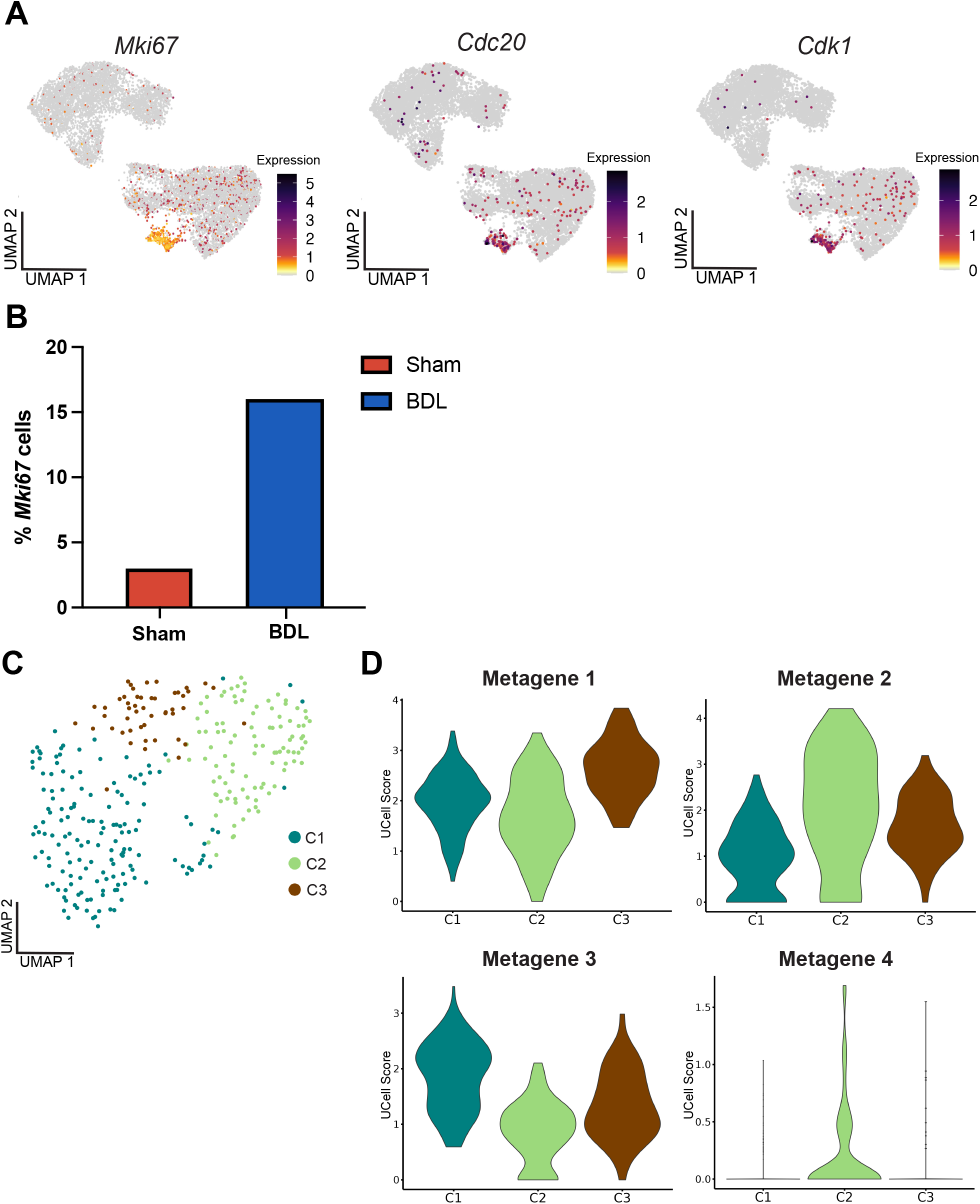
Metagene expression defines BEC heterogeneity in BDL-specific, highly proliferative BECs. (A) Cell cycle associated genes *Mki67, Cdc20*, and *Cdk1* are expressed throughout Sham and BDL subpopulations, but strongly upregulated in fifth subpopulation (BDL 5) found only following BDL. (B) The proportion of BECs expressing *Mki67* in Sham vs BDL increases >3-fold following injury, supporting significant proliferation induced by BDL. (C) Subclustering of BDL 5 reveals three distinct subclusters (C1-C3). (D) Differential enrichment of Metagene 1 in C3, Metagene 2 in C2, and Metagene 3 in C1 suggest that BEC-I, BEC-II, BEC-III upregulate cell cycle genes in response to BDL. Low expression of Metagene 4 across the same clusters suggests that BEC-IV are incapable of proliferating following injury or represent a transcriptomic program that is repressed following BDL.

### BEC metagene biomarkers and spatial neighborhoods are preserved in vitro

To explore *in vitro* BEC heterogeneity relative to metagenes, we isolated IHBD fragments from wild-type mice and cultured the cells to confluency. We then applied our 7-probe HCR FISH panel and quantified 1,578 cells in total (Figure 6A and Supplemental Table 8). BEC subpopulation proportions *in vitro* resembled those observed *in vivo*, with nominal differences (1% decrease in BEC-I; 3% decrease in BEC-II; 0.4% decrease in BEC-III; 4% increase in BEC-IV) (Figure 6B). Next, we asked if the spatial organization of BEC subpopulations observed *in vivo* is preserved in monolayer cultures *in vitro*. Qualitative analysis showed that cells of each BEC subpopulation localized near cells of the same subpopulation, forming homogeneous neighborhoods similar to those observed in vivo (Figure 6C). Together, these results demonstrate that metagene expression is preserved *in vitro* and continues to identify distinct subpopulations of BECs. Further, the spatial organization of BECs of the same subpopulation in discrete clusters in the absence of a mesenchymal niche may suggest that BEC heterogeneity is regulated in an epithelial-autonomous manner.

**Figure 6.**
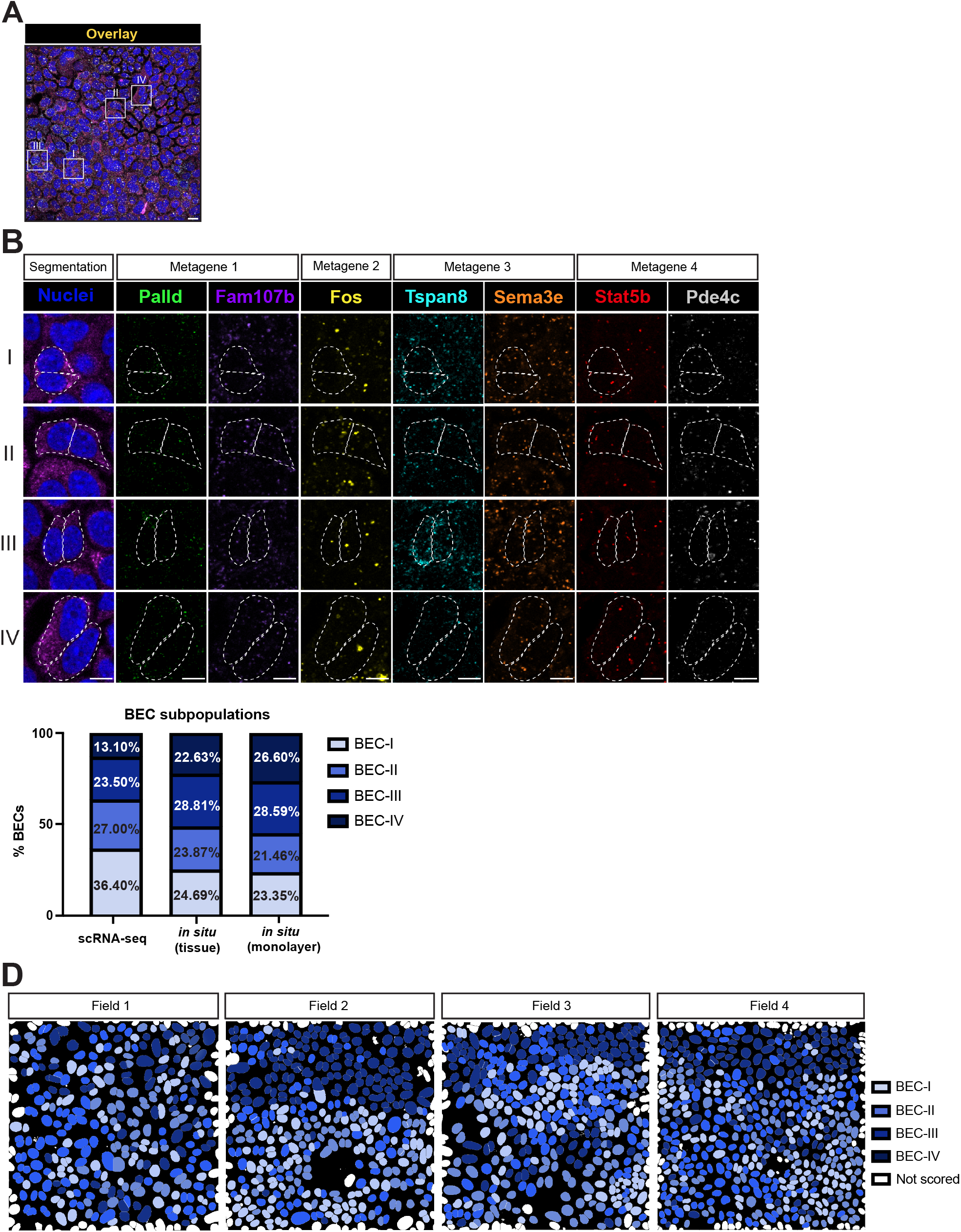
Metagene defined BEC heterogeneity is maintained in vitro. (A) Representative composite overlay of a 7-plex HCR-FISH imaging panel in BEC monolayers expanded for 7 days in vitro (white boxes indicate an area of metagene enrichment, shown at higher magnification in Figure 6B) (scale bar represents 20µm). (B) High-magnification single-channel images reveal metagene enrichment patterns that define BEC subpopulations (each row corresponds to a BEC subpopulation, and the dotted outlines labels the cells with the subpopulation identity) (scale bar represents 10µm). (C) The proportion of BEC subpopulations in BEC monolayers are similar to observed proportions in tissue and quantified from scRNA-seq. Minor shifts in relative abundance are observed across subpopulations but overall composition is largely preserved across *in vitro* model. (D) Spatial maps of the four imaging fields representing the 1,578 cells analyzed illustrate the organization of BEC subpopulations within monolayer cultures. Cells belonging to the same BEC subpopulation preferentially localize near one another, forming homogeneous spatial neighborhoods that recapitulate the spatial patterning observed *in vivo*. Data in panels (A–D) were generated from two independent monolayer cultures each established from a mouse liver, with two imaging fields per monolayer.

## DISCUSSION

Despite the power of scRNA-seq to define cellular heterogeneity across many tissue types and pathophysiological contexts, a clear consensus on intrahepatic BEC heterogeneity and subpopulation-specific biomarkers has remained elusive. Inherent technical challenges, including the relative scarcity of BECs versus other major hepatocellular lineages and the high transcriptomic similarities between BECs, have stymied these efforts ^9^. In this study, we first sought to identify BEC subpopulations and their associated biomarkers by applying a previously characterized Sox9EGFP allele. By FACS-purifying *Sox9*+ cells, we generated a scRNA-seq dataset focused specifically on BECs and HybHeps ^8^. Notably, we identified two subpopulations of HybHeps at homeostasis. These appear to adopt a more homogenous transcriptome – represented by a single population – following BDL. Though outside of the scope of the current manuscript, our results suggest a previously unappreciated degree of heterogeneity in *Sox9*+ peribiliary hepatocytes.

Because standard analysis of DEGs failed to identify cluster-specific BEC biomarkers, we pursued a metagene biomarker approach based on: (1) the co-expression of multiple genes in “on-target” clusters and (2) the de-enrichment of “off-target” metagenes in “on-target” clusters. Metagenes have been applied to represent transcriptome level patterns of gene expression via a limited number of gene biomarkers in microarray analysis and are seeing continued application to advanced multi-omics and spatial transcriptomics assays ^43^. We propose that metagenes can be used to identify four distinct subpopulations of BECs in mouse IHBDs. Applications of these metagenes could range from spatial localization of BECs, as in our HCR FISH assays, to identification of BEC subpopulations across experimental groups, as demonstrated in our Sham vs BDL analysis.

The most comprehensive hepatobiliary atlases to date have been generated from human liver tissue. These efforts have resulted in the identification of 6 BEC subpopulations (named Cholangiocyte (Chol)-1 through Chol-6), first in primary tissue by single nucleus RNA-seq (snRNA-seq) and more recently by scRNA-seq in organoids grown in refined media conditions that increase BEC heterogeneity *in vitro* ^10, 44^. Our results differ from human data in that we were unable to identify the previously described proliferative/progenitor-like and hepatocyte-like populations in homeostatic control mouse liver ^10^. Rather, we observe broad expression of hepatocyte genes *Hnf4a* and *Alb* across all BECs and a distinct proliferative cluster that emerges only after BDL. These results are unlikely to be due to under clustering, as our clustree analyses suggest that increasing clustering resolution to generate 6 Sham BEC subpopulations would be consistent with over clustering ^15^. Rather, our results resemble other scRNA-seq data of mouse BECs, where proliferative and hepatocyte-like clusters are reported in context of injury ^12, 13^. These data may suggest that the additional subpopulations in human atlases likely reflects biological differences and is possible that these represent fundamental differences between mouse and human biology. Alternatively, samples collected for human transcriptomic atlases may encompass a broader range of pathophysiological states, as human tissue acquisition for research is typically associated with underlying liver disease. It is notable that we initially identified 6 BEC clusters in our BDL samples, which we reduced to 5 by merging two clusters exhibiting similar enrichment of Metagene 3. Because computational identification of cellular subpopulations in scRNA-seq data analysis is susceptible to over-clustering, we prioritized a conservative approach that would also facilitate comparison of Sham and BDL subpopulations. Future studies may be able to further resolve BEC transcriptomic heterogeneity, ideally combined with lineage tracing to determine the cellular origin of additional BEC subpopulations in injury and disease models.

Another technical consideration is that our data set was generated using split-pool based scRNA-seq technology, enabling whole-transcript coverage ^45^. While *in situ* validation of candidate metagene biomarkers strongly supports the BEC subpopulations proposed here, whole transcript and 3’ end-counting scRNA-seq methods are known to yield subtle differences in transcript representation ^46^. Gene expression may also be an inherently poor readout of cell-cell heterogeneity in BECs. In this study and earlier work, we observe *Hnf4a* expression throughout BECs, but with little-to-no associated expression of HNF4A protein ^8^. Similarly, *Krt19* is enriched in a subpopulation of BECs while its protein product is ubiquitously expressed across all intrahepatic bile ducts with no apparent cell-cell difference in level ^8^. On one hand, potential discrepancies between BEC gene and protein expression could complicate efforts to easily identify subpopulation-specific protein biomarkers using transcriptomic data. However, it remains possible that protein expression patterns are more subpopulation-specific that gene expression patterns in BECs. While single cell proteomics remains an emerging field, substantial recent advances may soon facilitate its broader application ^47, 48^. Future efforts integrating multiple data sets and methodologies may hold potential to yield improved BEC subpopulation biomarkers.

An open question in the field is whether BECs even exist as well-defined and “fixed” subpopulations. Evidence from prior scRNA-seq studies comparing regional heterogeneity across the biliary tract supports a model where BECs are instead defined by highly plastic transcriptomic “states”. While regional differences between primary gallbladder, EHBD, and IHBD epithelium was evident by scRNA-seq, BECs isolated from three-dimensional organoids of the same regions demonstrated more homogenous transcriptomes ^49, 50^. More recent studies have shown that conventional human BEC organoid cultures are similarly homogenous at the single cell level, but that refined culture conditions can restore transcriptomic heterogeneity observed *in vivo* ^44^. This suggests that BEC heterogeneity *in vitro* is dependent on the proper milieu of extrinsic signaling present in culture conditions and is consistent with the known impact of culture media composition on cell fate in mouse and human intestinal organoids ^51, 52^. Our BDL results are also supportive of transcriptomic plasticity in BEC subpopulations, in that subpopulation-specific genes are downregulated while upregulated genes are most likely to be shared across multiple BEC subpopulations. However, despite larger transcriptomic changes, metagene expression remains useful in discriminating BEC heterogeneity in BDL. Similarly, our *in vitro* data demonstrate that BEC heterogeneity relative to metagene enrichment is preserved. While a limitation of the present study is that the full extent of transcriptomic heterogeneity in these conditions remains unknown, our results suggest that BEC metagenes may prove useful for assessing cellular heterogeneity in different contexts where pursuing scRNA-seq is either impossible or undesirable.

Inferring the functional and mechanistic impact of BEC heterogeneity remains a significant challenge. With the lack of robust biomarkers for BEC subpopulations coupled with the notable transcriptomic plasticity of BECs across physiological contexts, efforts are complicated when defining discrete BEC identities. Rather than representing fixed cell types, BEC subpopulations may instead reflect dynamic transcriptional states that are shaped by local signaling environments. With our approach, metagenes offer a framework to define core BEC subpopulation identities despite ongoing transcriptional remodeling. By enabling consistent identification of BEC subpopulations across conditions, metagenes may facilitate optimization of culture conditions to enrich for specific BEC states, providing a tractable platform for future work focused towards subpopulation-specific functional assays.

## METHODS

### Data and code availability

High-throughput sequencing data are available in Gene Expression Omnibus (GEO) under the following accession number: GSE 319345

### Mice

Development of Sox9^EGFP^ mice is described in Gong et al ^53^. The *Egfp* transgene was maintained at heterozygosity on the C57Bl/6 background. Sox9^EGFP^ mice were phenotyped visually by EGFP signal in the base of the hair follicle in the tail clipping on an epifluorescent microscope. All experiments were performed on adult mice of both sexes between 6-14 weeks of age.

For surgical procedures, Emory veterinarian staff performed both bile duct ligation and sham surgery as the control as previously reported ^54^. Briefly, Sox9^EGFP^+ mice were anesthetized using 1%–3% isoflurane in 100% oxygen and subjected to ∼2cm midline laparotomy using sterile surgical scissors. For the mice receiving the BDL, the common bile duct was exposed by gentle lifting of liver and caudal movement of the gut by using a sterile swab moistened with sterile 0.9% sodium chloride. The common bile duct was dissected from the portal vein using microserrated forceps and ligated with nonabsorbable 5-0 polyester suture, with a second cranial ligation placed in the same manner to thoroughly occlude the duct. The peritoneal cavity was rinsed with sterile 0.9% sodium chloride. All surgeries concluded with the closing of the peritoneum and skin using separate running sutures (6-0 monofilament, nonabsorbable). Mice were allowed to recover on heating pads and monitored twice daily for 2 days after surgery and then once daily for the remainder of the study. No adverse effects or unexpected mortality were observed. Mice were euthanized 7d post-surgery and were processed for either histology (n=3 per condition) or dissociated for FACS (n=2 per condition) as described below. All animal procedures were reviewed and approved by the Institutional Animal Care and Use Committee (IACUC) as Emory and UNC.

### Perfusion and tissue processing for histology

Tissues were fixed by intracardiac perfusion with cold PBS followed immediately by cold 4% paraformaldehyde (PFA; Thermo Fisher; 41678-5000) using a peristaltic pump (Kamoer DIpump550). Whole livers were dissected out and dropped into cold 4% PFA and stored overnight at 4°C. The tissues were then transferred to 30% sucrose and stored overnight 4°C. Individual lobes were dissected and embedded into Tissue-Tek Optimal Cutting Temperature media (Sakura Finetek USA, Torrance, CA), frozen down on dry ice and stored at -80°C until use.

### Immunofluorescence on tissue sections

For conventional histology, the embedded tissues were cut into 10μm sections on a cryostat (Leica; CM1860) and dried at room temperature for three hours. Tissues were rehydrated by washing with PBS for at least three minutes, permeabilized in 0.3% triton X-100 (Sigma; T8787-250ML) in PBS for 20 minutes, and blocked in 5% normal donkey serum (NDS; Jackson Immuno; 017-000-121) for 45 minutes. For all slides, primary antibody was applied overnight at 4°C in PBS. The next day, primary antibody solution was discarded, and slides were rinsed 3×5min with PBS. Secondary antibody raised against host species of the primary antibody was applied in PBS for 1hr at RT. After staining, antibody solution was discarded and bisbenzimide (Sigma; 14530) was applied at 1:1000 in PBS for 10min to label nuclei. Slides were then washed 3×5min with PBS, mounted with ProLong Glass Antifade (Thermo Fisher; P36980), and allowed to cure overnight at RT in the dark. Images were acquired on a laser scanning Confocal Microscope (Nikon A1R HD25) in the Emory Integrated Cellular Imaging Core using the 60X oil objective (NA 1.4 and WD 0.14 mm).

#### Antibody usage for immunostaining

Primary antibodies were used at the following concentration for immunostaining: 2.5μg/mL anti-EpCAM/CD326 (BioLegend; 118201). Secondary antibodies were used for immunostaining: 4μg/mL donkey anti-rat Alexa Fluor 555 (ThermoFisher; A31572) and 4μg/mL donkey anti-rat Alexa Fluor 647 (ThermoFisher; A21208).

### Intrahepatic bile duct dissociation and single cell isolation for FACS

BECs were isolated as previously described, with minor modifications ^7^. Liver lobes were dissected, specifically avoiding extrahepatic duct and gallbladder to prevent contamination. Tissue was rinsed in DPBS and then placed in petri dish. For mechanical digestion, tissue was minced to small pieces (<0.5cm^2^) using a razor blade and transferred to 50mL conical by resuspending in prewarmed 37°C Liver Wash Buffer [LWB; DMEM-H (Fisher Scientific; 11965118), 1% fetal bovine serum (Genesee Scientific; 25-525H), 1% GlutaMAX (Fisher Scientific; 35050061), 1% Pen/Strip (Fisher Scientific; 15140122)]. Tissue was allowed to sediment and then supernatant was discarded by pipetting. Tissue was rinsed in fresh prewarmed LWB and then discarded after sedimentation 2 more times. The tissue was pelleted in a swinging bucket centrifuge at 4°C and 600g for 5min. Supernatant was discarded and sample was resuspended in RT Liver Digest Buffer [0.15 mg/mL collagenase type XI (Sigma Aldrich; C9407), 0.3 U/mL dispase (Corning; 354235), 200 ug/mL DNase (Sigma Aldrich; DN25)] and incubated at 37°C on an end-over-end rotator for 90min. Every 30min the sample was retrieved and resuspended by pipetting vigorously for one minute with a P1000 micropipette set to 1mL as an additional mechanical digestion. After 90min, an equivalent volume of LWB was added to the dissociation and the tissue was pelleted at 4°C and 200g for 5min. The supernatant was carefully decanted to ensure the pellet remained intact and then the sample was washed again with LWB, pelleted, and supernatant decanted. Samples were resuspended in 5mL RBC lysis buffer (BioLegend; 420301) diluted in molecular grade water and incubated on ice for 5min with periodic agitation. RBC Lysis was quenched using 15mL DPBS and the sample was pelleted at 4°C and 600g for 5min and supernatant decanted. For culturing BEC monolayers, dissociation was stopped at this point, and duct fragments were used for organoid culture.

For single cell isolation, samples were resuspended with 3mL TrypLE (Fisher Scientific; 12605010) + 10µM Y-27632 (Selleck Chemicals; S1049) and incubated at 37°C for 12min. Every 2min, the sample was pipetted vigorously for 1min with a P1000 micropipette set to 1mL, to mechanically dissociate IHBDs. After 12min, the TrypLE was quenched using 15mL DPBS, and the sample was pelleted at 4°C and 600g for 5min. The supernatant was decanted, and pellet resuspended in Sort Media [Advanced DMEM/F12 (ThermoFisher; 12634028) with 2% B27 supplement without vitamin A (Thermo Fisher; 12587010), 1% N2 supplement (Thermo Fisher; 17502048), 2mM Glutamax (Thermo Fisher; 35050061), 10mM HEPES (Thermo Fisher; 15630080), 100 U/mL Pen/Strip (Fisher Scientific; 15140122), and 0.1% DNase (Sigma-Aldrich; DN25)]. After thoroughly resuspending, sample was filtered through a 40µm cell strainer (Falcon; 352340). Single cells were stained with anti-CD31-APC (1:100) (BioLegend; 102510), anti-CD45-APC (1:100) (BioLegend; 103112), and anti-CD326-APC/Cy7 (1:100) (BioLegend; 118218) for 1hr on ice. Cells were rinsed in Sort Media, pelleted at 4°C and 600g for 5min, and supernatant decanted. Cells were resuspended in sort media. 5uL of 7-AAD-APC (BioLegend; 420404) and 5 μL Annexin V-APC (BioLegend; 640941) were added to resuspended cells to distinguish dead and dying cells, respectively. Cells were immediately analyzed and collected on a Sony SH800S cell sorter. Gating scheme was designed to discriminate against non-epithelial cells by excluding CD31+, CD45+, 7-AAD+ and Annexin V cells and selected based on EGFP+ signal (Supplemental Figure 1A-B).

### Single Cell RNA-sequencing

EGFP+ BECs isolated by FACS were fixed and processed for single cell RNA sequencing following manufacturer’s guidelines for whole cells (Parse Biosciences; SB1001). Libraries were prepared using split-pool barcoding defined by the Evercode WT V2 kit (Parse Biosciences; SB1001). Pooled libraries were sequenced on the Illumina NovaSeq S4 flow cell in one lane (R1: 74 cycles, i7 index: 6 cycles, R2: 86 cycles, i5 index: 0 cycles) in the Emory Nonhuman Primate Genomics Core.

#### scRNA-seq analysis

The Parse Biosciences bioinformatics pipeline (v0.9.6p) was used to align reads to the *Mus musculus* reference genome (RefSeq GRCm39). The resulting filtered feature count matrices were further processed and analyzed using the *Seurat* package (v5.0.1) ^55^. Low-quality cells were filtered out if they contained fewer than 200 unique molecular identifiers (UMIs) or fewer than 100 detected features. Seurat objects for individual samples were merged using the *merge()* function.

Data normalization was performed using Seurat’s default LogNormalize method via the *NormalizeData()* function. Highly variable features were identified using *FindVariableFeatures()*, and data scaling was carried out using the *ScaleData()* function. Principal component analysis (PCA) was performed with *RunPCA()*, and clustering was conducted using the *FindNeighbors()* and *FindClusters()* functions with the first 23 principal components and a clustering resolution of 0.6. The optimal clustering resolution was determined by evaluating the clustering tree generated with the *clustree* package (v0.5.1). Nonlinear dimensionality reduction was performed using uniform manifold approximation and projection (UMAP) via the *RunUMAP()* function implemented in Seurat.

Cluster-specific marker genes and genes differentially expressed between conditions were identified through differential gene expression (DGE) analysis using the *FindAllMarkers()* and *FindMarkers()* functions, with the MAST statistical test (Finak et al., 2015) implemented in Seurat (Supplemental Tables 1, 2, and 4-7).

Single-sample gene set enrichment analysis (ssGSEA) was performed using the *runEscape()* function from the *escape* package (v2.2.3). For differential enrichment analysis, ssGSEA scores were first normalized using the *performNormalization()* function. The normalized scores were then compared across clusters using the *FindAllMarkers()* function with the Wilcoxon rank-sum test. Additionally, gene signature scores were calculated using *UCell* (v2.10.1).

For gene set enrichment analysis (GSEA), the DGE results table was filtered to include only genes expressed in at least 30% of the cells (pct.1 > 0.3) from the BDL samples. GSEA was performed using the GSEA() function from the clusterProfiler package (v4.14.6).

#### Metagenes

Sham-specific clusters (Sham 1, Sham 2, Sham 3, and Sham 4) were subset and converted to the H5AD file format using the *SeuratDisk* package (v0.0.0.9015). The minimal combinations of marker genes sufficient to distinguish among the Sham clusters were identified using NS-Forest (v3.9) with the following parameters: *n_trees = 1000, n_top_genes = 30, n_binary_genes = 25, n_genes_eval = 20, and beta = 0*.*5*.

UCell scores were calculated for each set of cluster-specific genes (metagenes) identified by NS-Forest, and the resulting metagene signature scores were visualized using the *VlnPlot()* and *FeaturePlot()* functions.

### Mouse intrahepatic biliary organoid and monolayer culture

#### Whole duct culture

Following the intrahepatic bile duct dissociation, ductal prep was resuspended in 500µL-1mL Advanced DMEM/F12 (Gibco) based on pellet size. 10µL aliquots from the ductal prep was examined by light microscopy to qualitatively determine cellular density for plating. 5-10µL of the ductal prep was further diluted in Advanced DMEM/F12 (Gibco), and Cultrex Type II Growth Factor Reduced extracellular matrix (R&D Biosystems, Minneapolis, MN; 3533-010-02) was added to a final concentration of 70% Cultrex. Cultrex-duct suspensions were plated as 30 µL droplets per well in pre-warmed 48-well plates and allowed to polymerize at 37°C for 25 minutes. After polymerization, droplets were overlaid with 200mL Biliary Isolation Media [50% Advanced DMEM/F12, 40% WNT3A-conditioned media, 10% RSPO1-conditioned media, 2% B27 supplement without vitamin A, 1% N2 supplement, 2mM Glutamax, 10mM HEPES, 100 U/mL Pen/Strip, 50ng/mL recombinant murine EGF (Thermo Fisher; PMG8041), 100ng/mL recombinant human Noggin (VWR International; 120-10C), 100ng/mL recombinant human FGF10 (Peprotech; 100-26), 10nM recombinant human gastrin (Sigma; G9145), 5ng/mL recombinant human HGF (PeproTech; 100-39H), 10mM nicotinamide (Sigma Aldrich; N0636-100G), 10µM forskolin (Selleck Chemicals; S2449), and 10µM Y-27632]. Media were replaced every other day with Biliary Expansion Media (WNT3A-conditioned medium was removed and replaced with Advanced DMEM/F12, and Noggin and Y27632 were withdrawn from culture).

Organoids were cultured for 7 days between passages. For passaging, Cultrex droplets were mechanically dissociated by pipetting 1mL of 1% BSA in PBS in each well and resuspending the droplet, scrapping the bottom of the well to ensure full recovery. Organoids were pelleted in a swinging bucket centrifuge at 4°C and 600g for 5min. Supernatant was carefully discarded by pipetting and organoid pellet was resuspended in 1mL TrypLE (Fisher Scientific; 12605010) + 10µM Y-27632 (Selleck Chemicals; S1049) and incubated at 37°C for four minutes. TrypLE was quenched using 1mL of 1% BSA in PBS, the organoids were further dissociated by pipetting 25 times, and then the sample was pelleted at 4°C and 600g for 5min. Organoid fragments were re-plated at 1:2 ratio as 30 µL droplets (70% Cultrex: 30% Advanced DMEM/F12) in pre-warmed 48-well plates, as described above. After each passage, BECs were initially grown in Biliary

Isolation Media with WNT3A, Noggin, and Y27632 and switched to Biliary Expansion Media after the first media change 2 days later.

#### BEC monolayer culture

Following the second passage, BEC organoid fragments were passaged into chamber slides coated with 10% Cultrex. BECs were initially grown in Biliary Isolation Media with WNT3A, Noggin, and Y27632 and switched to BEC Expansion Media after the first media change 2 days later. Monolayers were expanded until confluent (7-10 days).

### 7-plex Hybridization Chain Reaction (HCR) FISH and immunofluorescence

HCR FISH was performed on both tissue sections and cultured BEC monolayers from wildtype mice (Sox9^EGFP^-) following manufacturer’s protocol for fixed-frozen tissue sections with some modification. For BEC monolayers, after media was removed from each well and the chamber device was removed, samples were treated identically to fixed-frozen tissue.

#### HCR FISH

Samples were rehydrated with PBS for at least 3min at room temperature. Samples were fixed by adding slides to slide mailers with 4% PFA made from 16% Paraformaldehyde stocks (PFA Thermo; 043368-9M) for 10min on ice. Slides were transferred into a slide mailer containing 50% EtOH for 5 min at room temperature. This was followed by sequential washes in 70% EtOH and then two washes in 100% EtOH. Slides were quickly rinsed with DPBS and extra liquid was removed using a Kimwipe. Slides were transferred to a prewarmed (37°C) humidifier slide box. Prewarmed (37°C) HCR Hybridization Buffer (Molecular Instruments, v3.0) was added to each slide and incubated for 15min at 37°C. Slides were then tapped on the slide of the box to remove the buffer and probes were added to prewarmed (37°C) Hybridization Buffer at either 16nM (*Palld, Stat5b, Pdde4c, Fos*) or at 8nM (*Sema3e, Fam107b, Tspan8*). Slides were covered with parafilm to prevent evaporation, molecular grade water was added to the humidifier box and slides were incubated for 16-18hrs at 37°C.

Parafilm was removed from each slide and excess buffer was carefully tapped off. Slides were quickly rinsed with prewarmed (37°C) HCR Probe Wash Buffer (Molecular Instruments, v3.0) and then transferred to a slide mailer. Prewarmed Probe Wash buffer was diluted with Saline-Sodium Citrate (SSC) (Invitrogen, 15557036) Tween (Thermo, 85113) buffer (75%:25%, 50%:50%, 25%:75%, 0%:100%) and slides were incubated in each dilution in slide mailers for 15 min at 37°C. Slides were then incubated with fresh 100% SSC-Tween for 5min at RT followed by a quick DPBS rinse. Excess liquid was dried with a Kimwipe and slides were transferred to a slide box. HCR Amplification Buffer (Molecular Instruments, v3.0) was prewarmed (37°C) for resuspension before adding to each slide and incubated for 30 min at RT. Hairpins were snap-cooled (3min incubation at 95C followed by 30 min at RT in dark drawer) and added to HCR Amplification Buffer at 60 nM. Slides were tapped to remove excess liquid and the hairpins diluted in HCR Amplification Buffer was added to each slide. Parafilm was added to prevent evaporation and slides were incubated 16-18hr at RT.

Parafilm was removed, and slides were incubated in SSC-Tween for 5 min at RT. Slides were then transferred to a slide mailer and incubated in fresh SSC-Tween for 30 min at RT; this step was repeated. Slides were then transferred back to a slide box for immunofluorescence.

#### Immunofluorescence

Tissues are washed 3×5min in PBS in slide mailer. Slides were transferred to staining box and tissues were blocked in 5% normal donkey serum (NDS; Jackson Immuno; 017-000-121) for 45 minutes and primary antibody was applied overnight at 4°C in PBS. The next day, primary antibody solution was discarded, and slides were rinsed 3×5min with PBS. Secondary antibody raised against host species of the primary antibody was applied in PBS overnight at 4°C in PBS. The next day, antibody solution was discarded and bisbenzimide (Sigma; 14530) was applied at 1:1000 in PBS for 10min to label nuclei. Slides were then washed 3×5min with PBS, mounted with ProLong Glass Antifade (Thermo Fisher; P36980), and allowed to cure overnight at RT in the dark.

### 9-color image acquisition on confocal microscope

Tissue samples and fixed monolayers were imaged on the Leica STELLARIS 8 in the Emory Integrated Cellular Imaging Core. Each laser (excitation, ex), emission range (em), and detector group are as follows: bisbenzimide [(405 ex, 425-450 em, HyDS1), *Palld*-488 (488 ex, 495-505 em, HyDX1), *Stat5b*-514 (520 ex, 540-555 em, HyDS2), *Pde4c-*546 (560 ex, 575-585 em, HyDS1), *Sema3e*-594 (594 ex, 605-625 em, HyDS3), CD326-647 (653 ex, 660-6-85 em, HyDS3), *Fos*-700(700 ex, 705-740 em, HyDX2), *Fam107b*-750 (750 ex, 755-770 em, HyDX1), and *Tspan8*-790 (790 ex, 800-850 em, HyDX2)]. Channels were acquired in pairs (except 405) simultaneously as listed: [(bisbenzimide (Hoechst 33342, ThermoFisher, 62249), (*Palld* and CD326), (Stat5b and Fos), (Pde4c and Fam107b), and (Sema3e and Tspan8)]. Each pair was acquired as an entire frame before switching to the next pair. All image fields from tissue samples were acquired using the 63X oil objective (HC PL APO 63x/1.40 NA oil CS2) using a range of 1.49-5.38x zoom and a z-step no larger than 0.3µm. All image fields from fixed monolayers were acquired using the 40X oil objective (HC PL APO 40x/1.3 NA oil, PH3 CS2) with a z-step no larger than 0.3µm.

### 9-color image quantification and data analysis

Tissue samples and fixed monolayers were quantified using the Leica Application Suite (LAS) X (RRID:SCR_013673). Maximum Intensity Projection was performed on each z-stack image. For tissue sections, segmentation of BECs was based on positive CD326 signal, resulting in an ROI for each BEC imaged. Because the CD326 stain failed on BEC monolayers, segmentation of BECs was based on autofluorescence in the 647 channel. Mean intensity was calculated in the LAS X software for each individual channel (Supplemental Table 3 and 4).

#### Metagene Scoring

The mean intensity values for each channel corresponding to the 7 metagenes (488-*Palld*, 514-*Stat5b*, 546-*Pde4c*, 594-*Sema3e*, 700-*Fos*, 750-*Fam107b*, 790-*Tspan8*) was normalized by z-scoring in both tissue sections and monolayers. For tissue sections, z-scoring was calculated across all fields within each biological replicate (Supplemental Table 3). For monolayers, z-scoring was calculated within each field for each biological replicate (Supplemental Table 8). Using the z-scores, a metagene score was calculated for each cell based on the average z-score of the gene(s) composing the metagene (Supplemental Table 3 and 8). The identity for each BEC was determined by the highest metagene score (Supplemental Table 3 and 8).

#### Duct Diameter

The duct diameter was calculated by averaging the length of 2 sets of perpendicular lines (Supplemental Figure 5). Each line was drawn across the duct lumen, from the basolateral membranes of opposing BECs.

## FIGURE LEGENDS

**Supplementary Figure 1. *Transcriptomic analysis of Sox9EGFP+ cells in sham and BDL***. (A) Strategy for FACS isolation of Sox9^EGFP+^ cells from primary-isolated mouse liver tissue. (B) Final QC metrics show downstream analyses on cells with more than 1000 transcripts detected and with mitochondrial RNA content less than 50%. (C) Visualization of the clustree output, with nine different resolutions based on unsupervised clustering. Black arrow indicates the resolution for our clusters (res = 0.6; 16 total clusters). (D) Representative UMAP of the 16 clusters, labeled to correspond to the numbers in Supplemental Figure 1C. (E) Exclusion of cluster 11 is based on *Pecam1* expression which marks contaminating endothelial cells. (F) The four mouse samples cluster based on experimental condition, suggesting gene expression changes are a result of cholestasis and drive the major differences between clusters (Sham: n=2 mice and BDL: n= 2 mice).

**Supplementary Figure 2. *Gene signature scoring identifies BEC and HybHep clusters and reflects limited variability between BEC clusters***. (A) UMAP visualization of *Egfp* expression across all clusters. *Egfp* is enriched in Sham 1–4 and BDL 1–5 clusters, consistent with BEC identity, and reduced in Sham 5–6 and BDL 6 clusters, consistent with HybHep populations. (B– D) Violin plots showing UCell enrichment scores for transcriptomic signatures derived from previously published bulk RNA-seq of Sox9^EGFP^ populations. Gene signatures associated with GFP^high^ and GFP^low^ populations, which correspond to BECs, are enriched in Sham 1–4 and BDL 1–5 clusters. GFP^sub^ signature associated with HybHeps is enriched in Sham 5–6 and BDL 6 clusters. (E–J) Violin plots showing enrichment of published gene sets associated with signaling pathways and cellular features previously studied in BEC biology, including YAP, Notch, primary cilium, Hedgehog, extracellular matrix (ECM), and TGF-β signaling. These pathways show broad expression across BEC clusters with variable enrichment patterns between Sham and BDL conditions. Notably, Hedgeog signaling and ECM signatures are enriched in Sham 5–6 and BDL 6 clusters, suggesting a potential role in HybHeps. (K) Enrichment of a previously defined duct versus ductule gene signature suggests Sham 3 and consistent with BDL 3 are BECs localized to large ducts.

**Supplementary Figure 3. *Classical hepatocyte and BEC markers are broadly expressed across BEC clusters***. (A) Expression of canonical hepatocyte markers (*Hnf4a* and *Alb*) are detectable across multiple BEC clusters, with variable expression levels and no clear cluster-specific pattern. (B) Expression of reported small ductule–associated markers (*Hrh1, Nfatc2*, and *Sp1*) across clusters. *Hrh1* shows localized enrichment to a subpopulation of BECs within Sham 3 in Sham 3 and BDL 3 and 4. The remaining markers are broadly expressed among BEC clusters. (C) Expression of reported large bile duct–associated markers (*Slc10a2, Slc4a2, Sctr, Nfatc3, Mtnr1a*, and *Cftr*). Most markers exhibit widespread expression across BEC clusters.

**Supplementary Figure 4. *NSForest identifies minimal gene sets with binary expression defining four BEC subpopulations in sham***. Across sham BEC clusters, each metagene shows strong enrichment in a single cluster and minimal expression in the remaining sham clusters, achieving binary expression. **(A–B)** Expression of *Palld* and *Fam107b* (Metagene 1) defines the Sham 1 subpopulation. **(C)** Expression of *Fos* (Metagene 2) uniquely defines Sham 2. **(D–E)** Expression of *Tspan8* and *Sema3e* (Metagene 3) defines the Sham 3 subpopulation. **(F–G)** Expression of *Stat5b* and *Pde4c* (Metagene 4) defines the Sham 4 subpopulation.

**Supplementary Figure 5. *In situ metagene analysis of duct diameters reveals trends in BEC-III and BEC-IV to be associated with large and small ducts, respectively***. (A) Representative imaging field for duct quantification. Two sets of perpendicular lines are drawn across the ductal lumen from the basolateral membranes of opposing BECs to calculate the duct diameter. (B) Representative calculation for duct diameter (µm): the average of the length of the 4 lines drawn in Supplemental Figure 5A. (C) Quantification of BEC subpopulation composition across all IHBDs imaged by HCR-FISH. Each bar represents the proportion of a given BEC subpopulation within a single duct. All IHBDs analyzed contain at least two distinct BEC subpopulations, demonstrating transcriptional heterogeneity within individual ducts.

**Supplementary Figure 6. *Metagene enrichment matches BDL to Sham BEC subpopulations and reveals injury-induced transcriptional programs***. (A) Expression of **Metagenes 1, 2, and 4** across BDL BEC clusters demonstrates that each metagene is selectively enriched for a single BDL cluster. Because metagene 3 expression is enriched in both BDL 3 and 4, these clusters are both representing BEC-III and merge to one cluster (Figure 4B). (B) Differential gene expression analysis identifies a combination of shared and subpopulation-specific DEGs in response to BDL. In total, 293 genes are shared across at least two BEC subpopulation comparisons, alongside uniquely upregulated and downregulated genes within individual subpopulations. (C) Gene set enrichment analysis reveals shared pathways (cell adhesion, cell locomotion, and cell motility) across all four BEC subpopulations. (D) Downregulated DEGs from each pairwise comparison downregulated in BDL BEC clusters (BDL 1-4), validating the DE analysis.

**Supplementary Figure 7. *Clustree identifies three stable subclusters in the BDL 5 cluster***.

Black arrow indicates cluster resolution selected for analysis.

**Supplemental Table 1. *Differential expression analysis for Sham 1-6 and BDL 1-7 clusters***.

**Supplemental Table 2. *Differential expression analysis for Sham 1-4 clusters***.

**Supplemental Table 3. *Metagene scores for BECs in tissue sections***.

**Supplemental Table 4. *Differential expression analysis for BDL 1 vs Sham 1***.

**Supplemental Table 5. *Differential expression analysis for BDL 2 vs Sham 2***.

**Supplemental Table 6. *Differential expression analysis for BDL 3 vs Sham 3***.

**Supplemental Table 7. *Differential expression analysis for BDL 4 vs Sham 4***.

**Supplemental Table 8. *Metagene scores for BECs in monolayers***.

## Notes

This study was funded by the NIH/NIDDK under award numbers R01DK132653 (Gracz) and F31DK134199 (Hrncir), and by the National Science Foundation Graduate Research Fellowship under Grant No. 1937971 and 2439564 (Kanakanui). Research reported in this publication was supported in part by the Emory Integrated Genomics Core (EIGC) shared resource of Winship Cancer Institute of Emory University and NIH/NCI under award number P30CA138292, by the Emory University Emory Integrated Cellular Imaging Core Facility (RRID:SCR_023534) and NIH under Award Number S10 OD032320-01, the Emory NPRC Genomics Core is supported in part by NIH P51 OD011132.

The authors declare no conflicts of interest

https://www.ncbi.nlm.nih.gov/geo/query/acc.cgi?acc=GSE319345

### Competing Interest Statement

The authors have declared no competing interest.

