## Supplementary figures and images for "Multi-gene biomarkers reveal spatial organization and subpopulation-specific damage response in intrahepatic biliary epithelial cells"

### Supplemental Figure 1

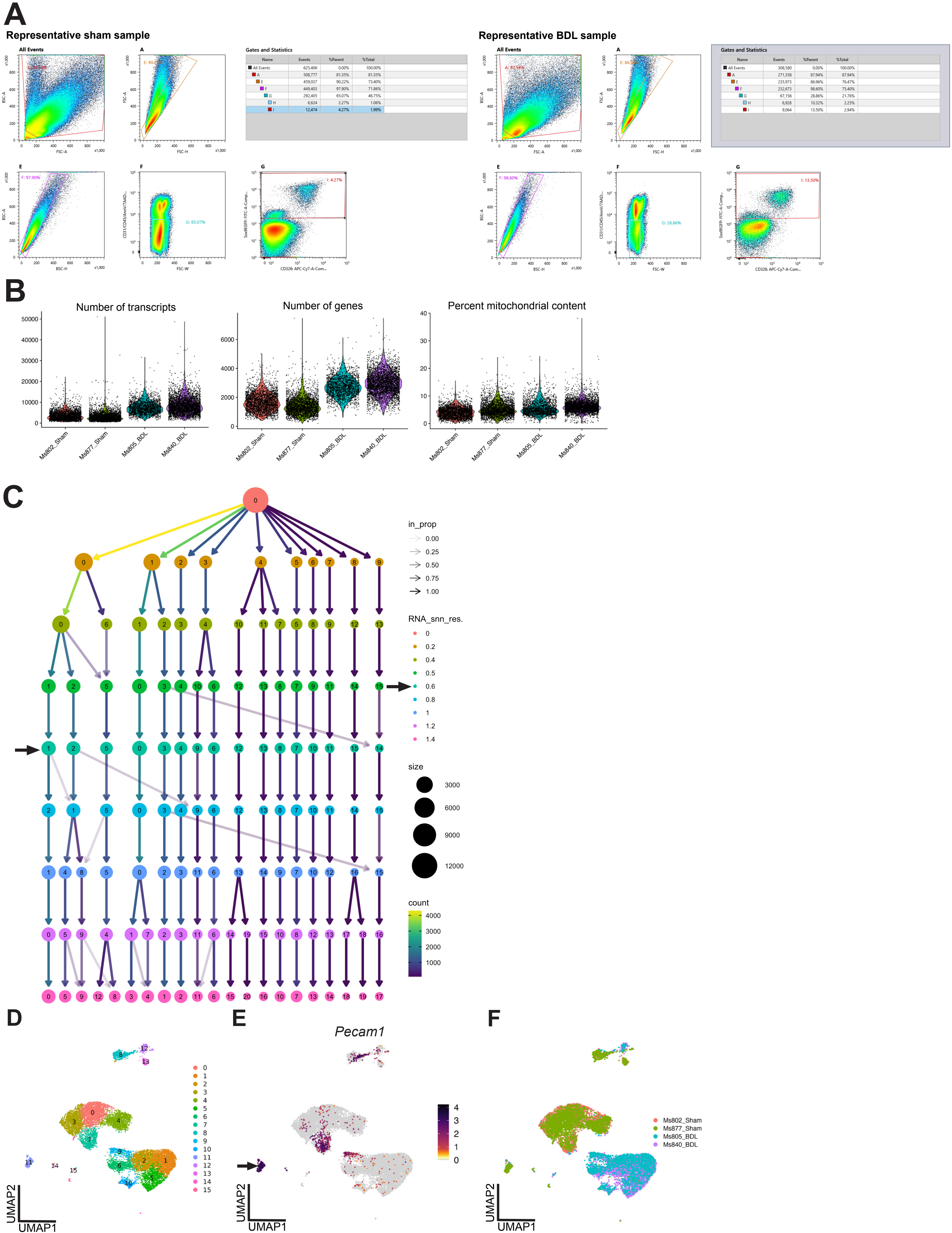

### Supplemental Figure 2

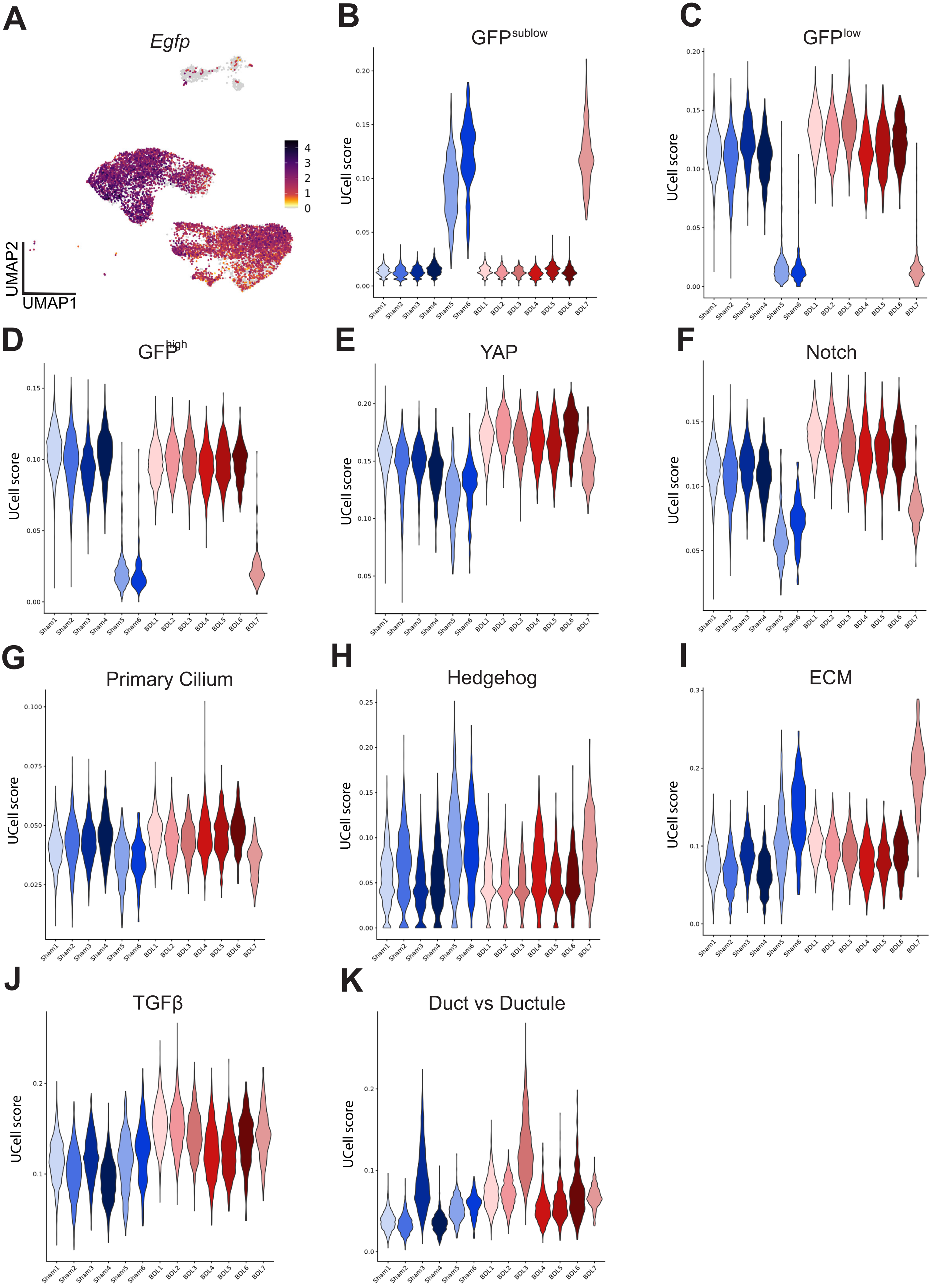

### Supplemental Figure 3

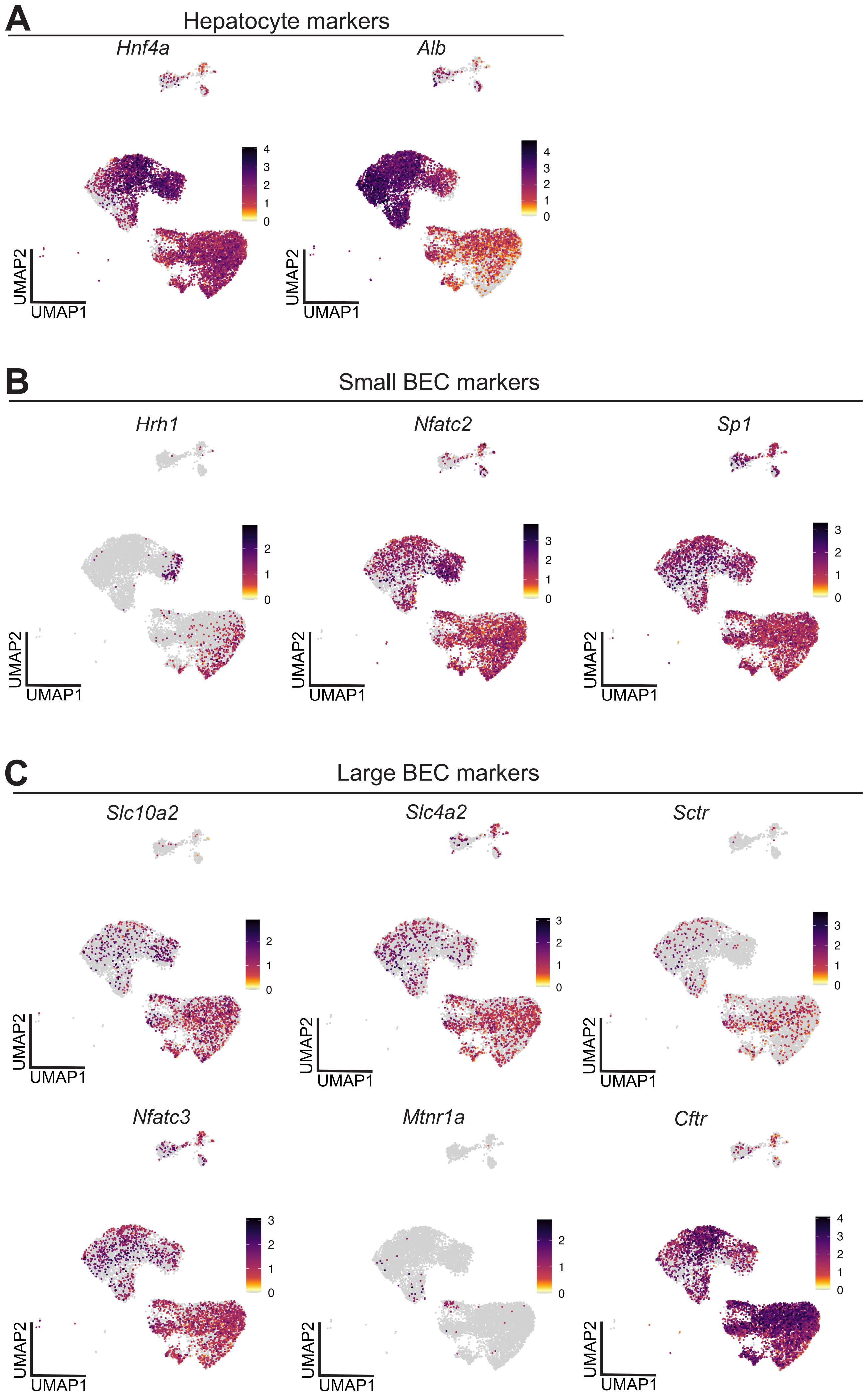

### Supplemental Figure 4

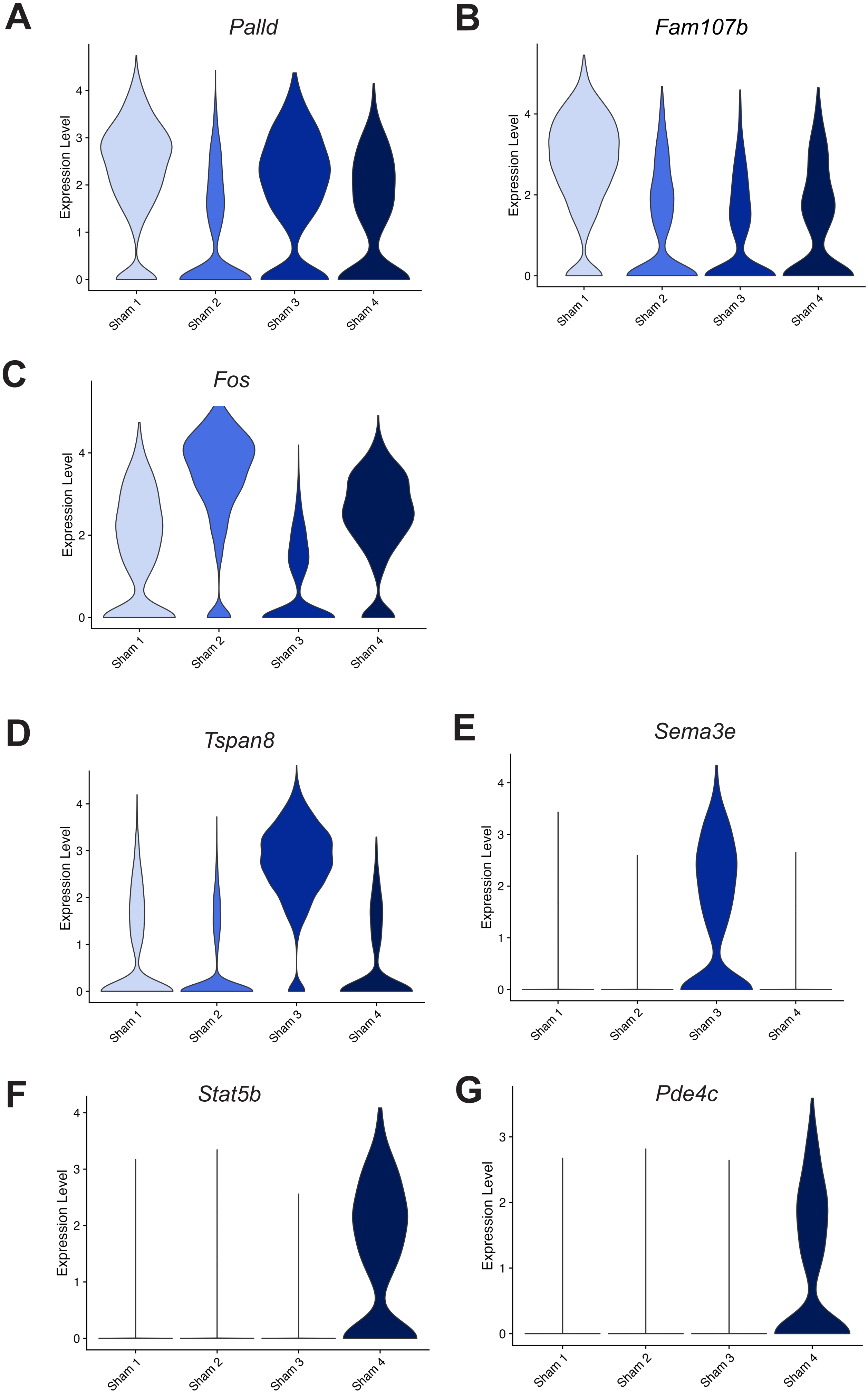

### Supplemental Figure 5

A

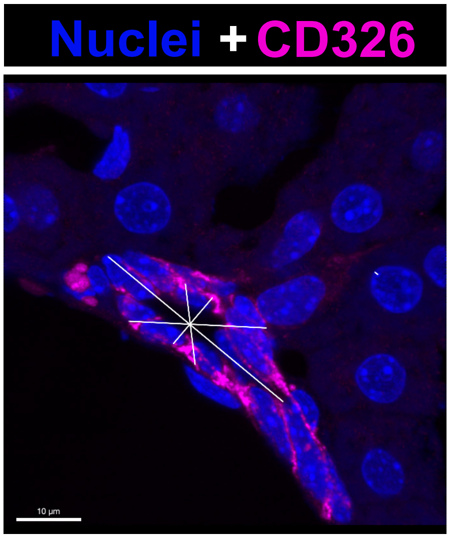


B

| | |
|---|---|
| Line 1 | 33.72 µm |
| Line 2 | 10.71 µm |
| Line 3 | 22.34 µm |
| Line 4 | 11.22 µm |
| AVERAGE | 19.50 µm |

C

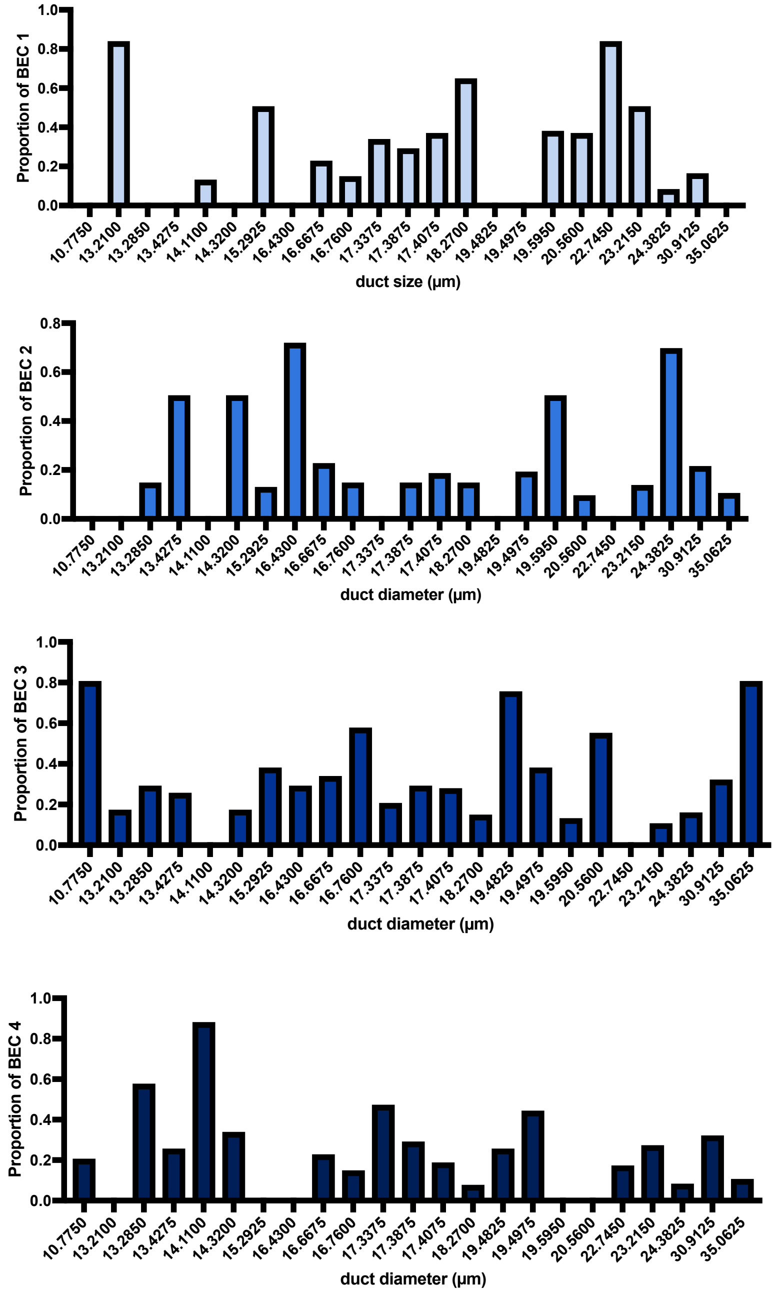

### Supplemental Figure 6

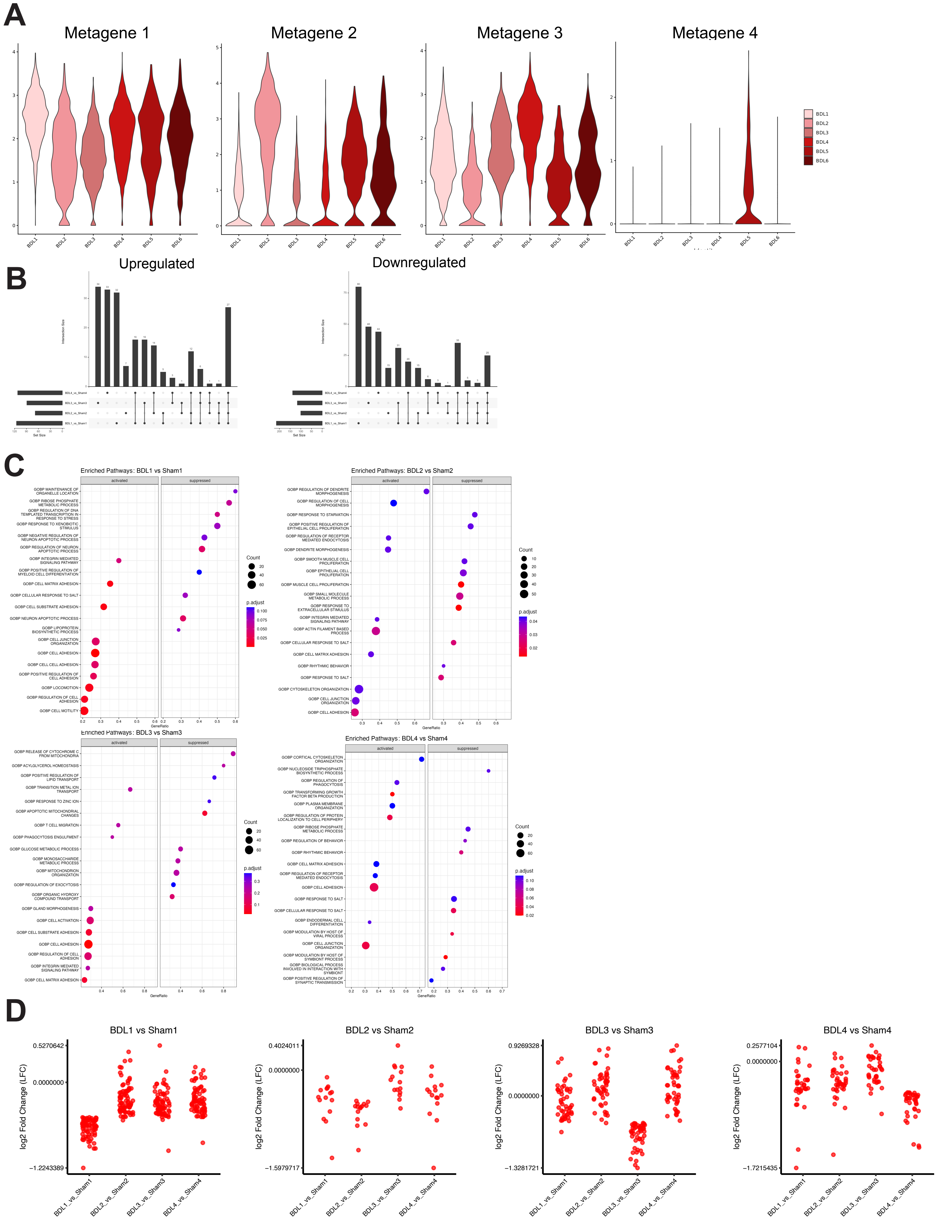

### Supplemental Figure 7

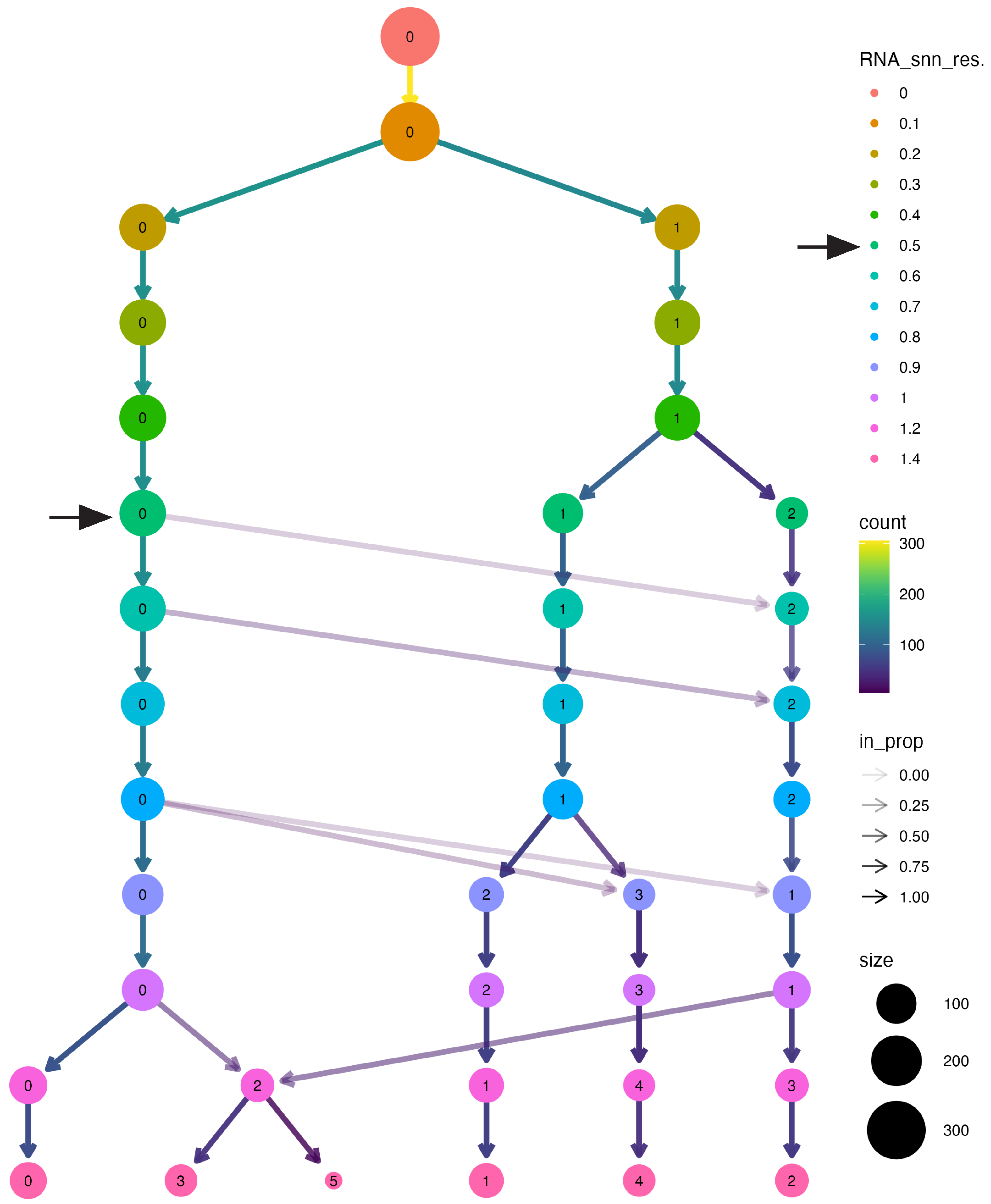
